# Peroxiredoxin 5 regulates osteogenic differentiation via interaction with hnRNPK during bone regeneration

**DOI:** 10.1101/2022.06.09.495435

**Authors:** Eunjin Cho, Xiangguo Che, Mary Jasmin Ang, Seongmin Cheon, Jinkyung Lee, Kwang Soo Kim, Chang Hoon Lee, Sang-Yeop Lee, Hee-Young Yang, Changjong Moon, Chungoo Park, Je-Yong Choi, Tae-Hoon Lee

**Author notes:** Correspondence and requests for materials should be addressed to T-H.L.

## Abstract

Peroxiredoxin 5 (Prdx5) is involved in pathophysiological regulation via the stress-induced cellular response. However, the function of Prdx5 in the bone remains largely unknown. Here, we show that Prdx5 is involved in osteoclast and osteoblast differentiation, resulting in osteoporotic phenotypes in *Prdx5* knockout (*Prdx5*^Ko^) mice. Through immunoprecipitation and liquid chromatography combined with tandem mass spectrometry analysis, heterogeneous nuclear ribonucleoprotein K (hnRNPK) was identified as a potential binding partner of Prdx5 during osteoblast differentiation *in vitro*. We found that Prdx5 acts as a negative regulator of hnRNPK-mediated osteocalcin (*Ocn*) expression. In addition, transcriptomic analysis revealed that *in vitro* differentiated osteoclasts from the bone marrow-derived macrophages of *Prdx5*^Ko^ mice showed enhanced expression of several osteoclast-related genes. These findings indicate that Prdx5 might contribute to the maintenance of bone homeostasis by regulating osteoblast differentiation. This study proposes a new function of Prdx5 in bone remodeling that may be used in developing therapeutic strategies for bone diseases.

## Introduction

The bone is remodeled through continuous replacement of old tissues by new tissues (Sims & Walsh, 2012). This process involves bone deposition or production by osteoblasts and bone resorption by osteoclasts, which are responsible for the breakdown of old bone tissues (Knothe Tate et al., 2004; Yang et al., 2020). Remodeling allows bones to adapt to stress; for instance, bones can become thick and strong when subjected to stress, and bones that are not exposed to regular stress begin to lose mass (Wang et al., 2022). However, the balance between osteoclasts and osteoblasts is critical in bone remodeling; an imbalance between these cells can lead to bone loss (Weitzmann & Ofotokun, 2016).

Peroxiredoxins (Prdxs) are a large superfamily of antioxidant enzymes that reduce peroxides (Rhee, 2016). Prdxs are classified as 1-Cys (Prdx1–5) and 2-Cys (Prdx6) based on their conserved cysteine residues (Seong et al., 2021), and they protect cells from oxidative stress (Lee et al., 2020; Rhee, 2016). Prdx6 inhibits bone formation in newborn mice (Park et al., 2019). Thioredoxin-1 induces osteoclast differentiation, which is suppressed by glutathione peroxidase-1 and Prdx1 (Lean et al., 2004). Prdx5 acts as a mitochondrial antioxidant and regulates ciliogenesis, adipogenesis, and fibrogenesis (Choi et al., 2019; Ji et al., 2019; Kim et al., 2018). Furthermore, it ameliorates obesity-induced non-alcoholic fatty liver disease by modulating mitochondrial reactive oxygen species (ROS) (Kim et al., 2020). From a biochemical perspective, Prdx5 regulates the activation of cyclin-dependent kinase 5 and Ca^2+^/calcineurin-Drp1, Jak2–Stat5 modulation during pathogenic conditions via antioxidant activity, and the protein–protein interactions (Choi et al., 2013; Chung et al., 2010; Park et al., 2016; Park et al., 2017; Yang et al., 2010). However, the role of Prdx5 in bone remodeling has not yet been studied.

Heterogeneous nuclear ribonucleoproteins (hnRNPs) are a family of nuclear proteins that function in mRNA biogenesis, including pre-mRNA splicing (Expert-Bezancon et al., 2002), transport of mRNA from the nucleus to the cytosol (Michael et al., 1997), and translation (Ostareck et al., 1997). Heterogeneous nuclear ribonucleoprotein K (hnRNPK) is a unique member of this family, as it preferentially binds single-stranded DNA, whereas other RNPs bind RNA (Siomi et al., 1994). hnRNPK is a multifunctional molecule, which can act both in the cytosol and nucleus (Krecic & Swanson, 1999; Mikula et al., 2010), and has been implicated in different cellular processes, including gene transcription (Michelotti et al., 1996; Tomonaga & Levens, 1996) and chromatin remodeling (Denisenko & Bomsztyk, 2002), in addition to the more typical functions of splicing and mRNA transport to the cytoplasm (Dreyfuss et al., 1993). hnRNPK mutation in humans causes a Kabuki-like syndrome with skeletal abnormalities and facial dysmorphism (Wang et al., 2020); acute myeloid leukemia patients show aberrant hnRNPK expression (Gallardo et al., 2015). hnRNPK deletion in mice is embryonically lethal, and haploinsufficiency results in developmental defects with skeletal disorders and post-natal death (Au et al., 2018; Dentici et al., 2018; Gallardo et al., 2015). hnRNPK acts as a transcription factor and regulates translation by binding to promoters. In cancer, hnRNPK binds to the promoter regions of *c-myc* and *c-src* to elevate their transcription or binds to their mRNAs to control translation (Naarmann et al., 2008; Perrotti & Neviani, 2007; Ritchie et al., 2003). In the bone, hnRNPK interacts with glycogen synthase kinase 3 beta to promote osteoclast differentiation (Fan et al., 2015). During osteoblast differentiation, hnRNPK binds to the promoter region of osteocalcin (*Ocn*) and represses its transcription (Stains et al., 2005). However, hnRNPK requires other interacting proteins to regulate gene expression, and the underlying mechanisms in the bone are largely unexplored.

Here, we examined Prdxs during osteoblast and osteoclast differentiation *in vitro*. Interestingly, Prdx5 expression was significantly altered during cell differentiation, i.e., it was upregulated during osteogenesis but suppressed during osteoclastogenesis. Therefore, we defined the role of Prdx5 in the bone using *Prdx5*-deficient (Ko) mice. In micro-computed tomography (micro-CT) analysis, *Prdx5*^Ko^ mice showed osteoporosis-like phenotypes with increased osteoclast and reduced osteoblast differentiation. *Prdx5*^Ko^ mice showed a delay in osteogenesis in the calvarial defect model. To determine the interacting partners of Prdx5 during osteogenesis, we performed liquid chromatography combined with tandem mass spectrometry (LC-MS/MS) analysis. Among the binding partners, hnRNPK colocalized with Prdx5 during osteoblast differentiation. We suggested that Prdx5 controls hnRNPK translocation to inhibit its binding to the *Ocn* promoter for osteoblast differentiation. RNA-sequencing (RNA-seq) analysis showed a significant increase in the expression of osteoclast-related genes in the osteoclasts differentiated from bone marrow-derived macrophages (BMMs) of *Prdx5*^Ko^ mice compared to that in the osteoclasts of wild-type (WT) mice. These results indicate a new role of Prdx5 in bone biology, i.e., Prdx5 homeostasis is critical in bone remodeling. Therefore, Prdx5 may be useful for understanding and preventing osteoporotic diseases involving osteoclast activity.

## Results

### Prdx5 is controlled during bone cell differentiation

To elucidate whether Prdxs function in bone remodeling, we characterized the expression of all Prdxs (Prdx1–6) during osteoclast and osteoblast differentiation *in vitro* (Figure 1). First, calvaria-derived pre-osteoblasts were differentiated into osteoblasts via stimulation with bone morphogenic protein 2 (BMP2). The mRNA levels of *Prdx3* and *Prdx5* were significantly elevated upon BMP2 stimulation (Figure 1A). During osteoblast differentiation, Prdx2 and Prdx5 expression was increased by BMP2 stimulation (Figure 1B). To explore the function of Prdxs in osteoclast differentiation, BMMs were differentiated into osteoclasts via treatment with receptor activator of the nuclear factor-κB ligand (RANKL). The mRNA levels of *Prdx1* and *Prdx5* were significantly reduced (Figure 1C). Prdx4 and Prdx5 expression was altered by RANKL treatment (Figure 1D). Interestingly, Prdx5 levels were reduced during osteoclastogenesis but increased during osteogenesis, with a correlation between mRNA and protein expression. Therefore, we focused on Prdx5 as a potential regulator of bone remodeling.

**Figure 1.**
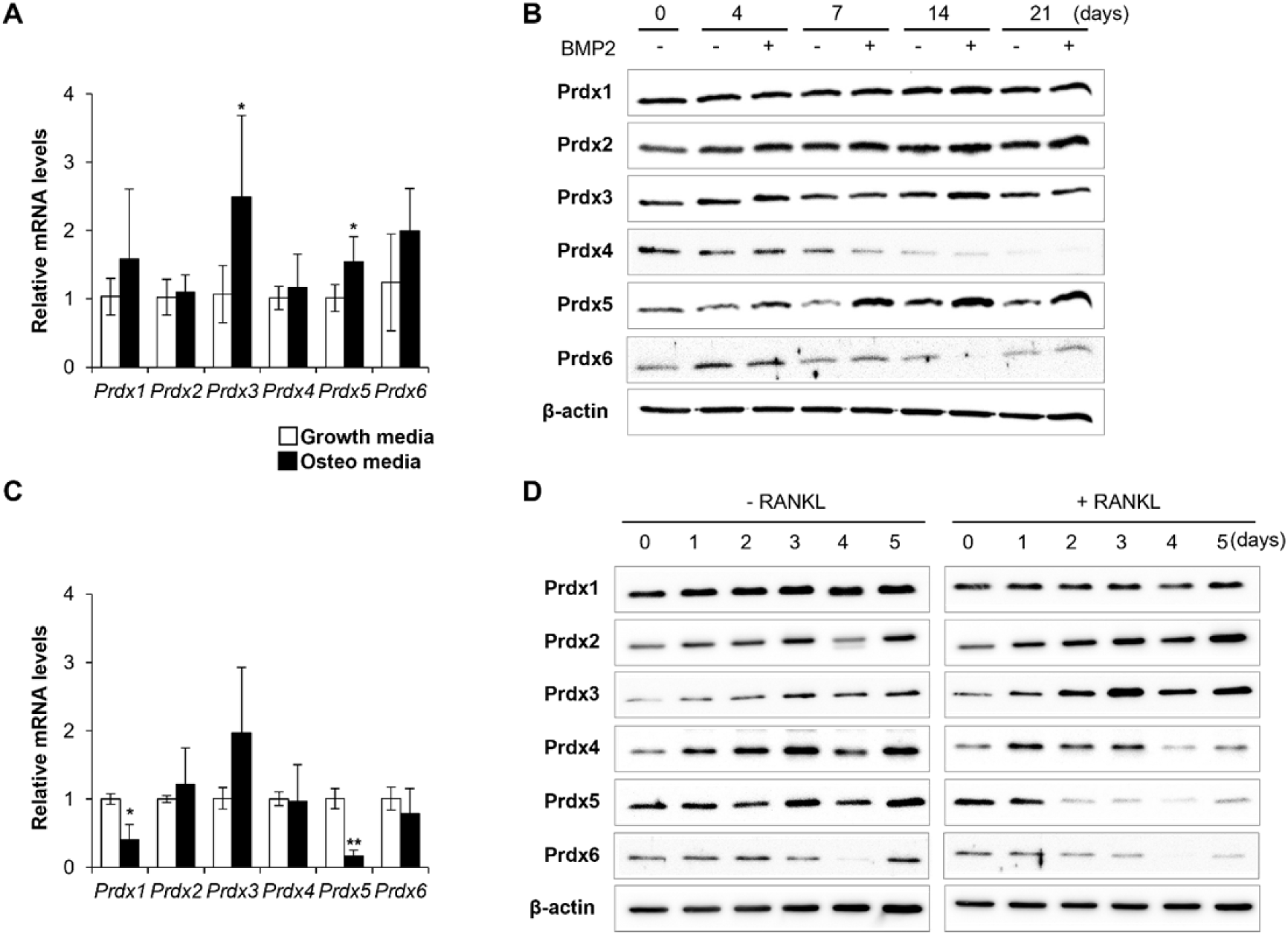
Prdx 5 expression is controlled during bone cell differentiation. (A) mRNA expression of Prdxs was determined, using qRT-PCR, in osteoblasts on day 7 after BMP2 stimulation. (B) Protein levels of Prdxs in osteoblasts were determined via western blotting. (C) mRNA levels of Prdxs were determined in osteoclasts on day 3 after RANKL stimulation. (D) Protein levels of Prdxs in osteoclasts were determined using western blotting. Graph depicts mean ± SD. \**p* < 0.05, \*\**p* < 0.01 via an unpaired two-tailed Student’s *t*-test.

### Abnormal expression of Prdx5 modulates osteoblastogenesis and osteoclastogenesis *in vitro*

To clarify the roles of Prdx5 in osteoblast and osteoclast differentiation, we thoroughly examined its expression *in vitro*. *Prdx5* mRNA expression was elevated by BMP2 stimulation on days 4 and 7 and decreased on day 14 (Figure 2A). However, Prdx expression was continuously upregulated till day 14. We isolated the precursor cells from *Prdx5*^Ko^ mice and examined osteoblast differentiation using alkaline phosphatase (ALP) staining. Osteoblast differentiation was strongly inhibited in *Prdx5*^Ko^ cells on day 7 (Figure 2B). To examine the expression of osteoblast-specific genes, we performed quantitative reverse transcription-PCR (qRT-PCR) on day 7 after BMP2 administration (Figure 2C). The mRNA levels of Runt-related transcription factor 2 (*Runx2*), *Alp*, and *Ocn* were increased. However, the upregulation in *Prdx5*^Ko^ cells was not significant compared to that in WT. These results suggest that Prdx5 is necessary for osteoblast differentiation, as it regulates osteogenic marker gene expression.

**Figure 2.**
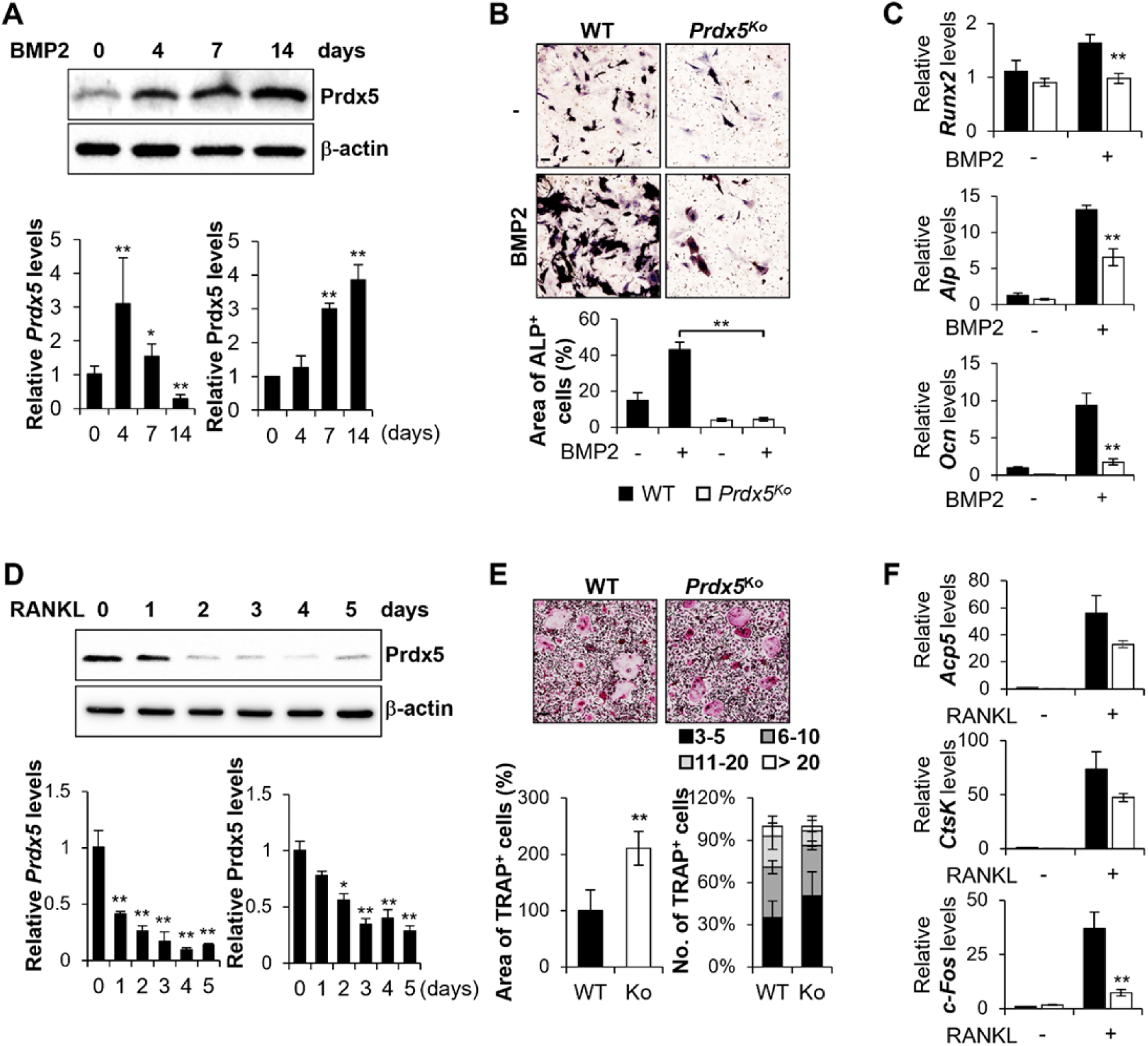
Abnormal expression of Prdx5 modulates osteoblastogenesis and osteoclastogenesis *in vitro*. (A, B, C) Mouse calvaria-derived pre-osteoblasts were differentiated into osteoblasts through BMP2 stimulation for indicated time periods. (A) Western blotting and qRT-PCR were performed to determine Prdx5 expression during osteoblastogenesis. (B) Pre-osteoblasts were isolated from WT and *Prdx5*^Ko^ mice and then differentiated into osteoblasts for 7 days. ALP staining was performed to determine the number of osteoblasts, and the area of ALP-positive cells was measured using the Image J software. (C) qRT-PCR was performed to determine osteogenic gene expression on day 7. (D, E, F) BMMs were differentiated into osteoclasts through 30 ng/mL M-CSF and 50 ng/mL RANKL stimulation for indicated time periods. (D) Western blotting and qRT-PCR were performed to determine Prdx5 expression during osteoclastogenesis. (E) BMMs were isolated from WT and *Prdx5*^Ko^ mice and then differentiated into osteoclasts for 4 days. TRAP staining was performed to determine the number of osteoclasts, and the area of TRAP- positive cells was measured. The number of multinucleated cells harboring the indicated nuclei was counted. (F) qRT-PCR was performed to determine the expression of osteoclast-related genes. Graph depicts mean ± sem. \**p* < 0.05, \*\**p* < 0.01 via an unpaired two-tailed Student’s *t*-test compared to control (0) or WT.

Next, we analyzed Prdx5 expression during osteoclast differentiation after RANKL and macrophage colony-stimulating factor (M-CSF) administration (Figure 2D–F). Prdx5 expression decreased from the first day of osteoclastogenesis (Figure 2D). The efficacy of osteoclast differentiation was examined in the BMMs from *Prdx5*^Ko^ mice (Figure 2E). To determine the number of differentiated osteoclasts, tartrate-resistant acid phosphatase (TRAP, *Acp5*) staining was performed. Interestingly, the *Prdx5*^Ko^ BMMs showed a 2-fold increase in TRAP-positive areas compared to the WT. Osteoclasts become multinucleated giant cells via cell–cell fusion to acquire bone resorption activity (Kodama & Kaito, 2020). Therefore, we measured the number of nuclei in a TRAP-positive cell as an indicator of cell fusion. *Prdx5*^Ko^ cells showed a smaller number of nuclei than WT. During osteoclastogenesis, the levels of *Acp5* and cathepsin K (*CtsK*) increase remarkably in mature osteoclasts, and the transcription factor c-Fos regulates nuclear factor of activated T cells cytoplasmic 1 (NFATc1)-mediated signaling pathways (Nagy & Penninger, 2015; Yang & Karsenty, 2002). The mRNA levels of *Acp5* and *CtsK* were significantly reduced in *Prdx5*^Ko^ cells on day 3 (Figure 2F) but increased up to the levels in WT on days 4 and 5 (Figure 2–figure supplement 1). These data suggest that, in *Prdx5*^Ko^, BMMs develop osteoclasts at a slower rate than that in WT. These differences are not altered at the maturation stage of osteoclasts *in vitro*.

To determine whether the osteoporosis phenotype in *Prdx5*^Ko^ mice is because of an increase in ROS levels, we determined ROS levels in cultured osteoblasts and osteoclasts from *Prdx5*^Ko^ and WT mice (Figure 2–figure supplement 2). In osteoblasts, the precursor cells from *Prdx5*^Ko^ mice showed slightly reduced ROS levels than those from WT. In osteoblasts, ROS levels increased in WT cells after BMP2 stimulation, whereas ROS in *Prdx5*^Ko^ cells were maintained at the precursor levels. However, ROS levels were not changed upon RANKL-stimulation during osteoclast differentiation in WT and *Prdx5*^Ko^ cells. These data suggest that ROS are not significantly involved in Prdx5-mediated osteoblast or osteoclast differentiation. Therefore, Prdx5 may regulate alternative mechanisms of bone cell differentiation, apart from acting as an antioxidant.

### *Prdx5*^Ko^ mice show enhanced osteoporotic phenotypes

To determine the role of Prdx5 in bone remodeling *in vivo*, we analyzed bone parameters in *Prdx5*^Ko^ mice (Figure 3). Micro-CT analysis of the distal femurs showed that *Prdx5*^Ko^ mice had low bone mineral density (BMD) and trabecular number (Tb. N) and an increased trabecular bone space (Tb. Sp) compared to those in their WT littermates (Figure 3B). Additionally, *Prdx5*^Ko^ mice showed reduced trabecular volume (Tb. V) and thickness (Tb. Th), which suggested reduced trabecular bone formation in *Prdx5*^Ko^ mice compared to that in WT. To determine bone-related cytokine levels in the serum, RANKL, osteoprotegerin (OPG), and BMP2 levels were examined (Figure 3C). In *Prdx5*^Ko^ mice, RANKL and OPG levels increased by 1.5-fold compared to those in WT. However, BMP2 levels were not altered in *Prdx5*^Ko^ mice. These findings suggested that osteoporosis-like phenotypes in *Prdx5*^Ko^ mice were mediated by an increase in RANKL expression.

**Figure 3.**
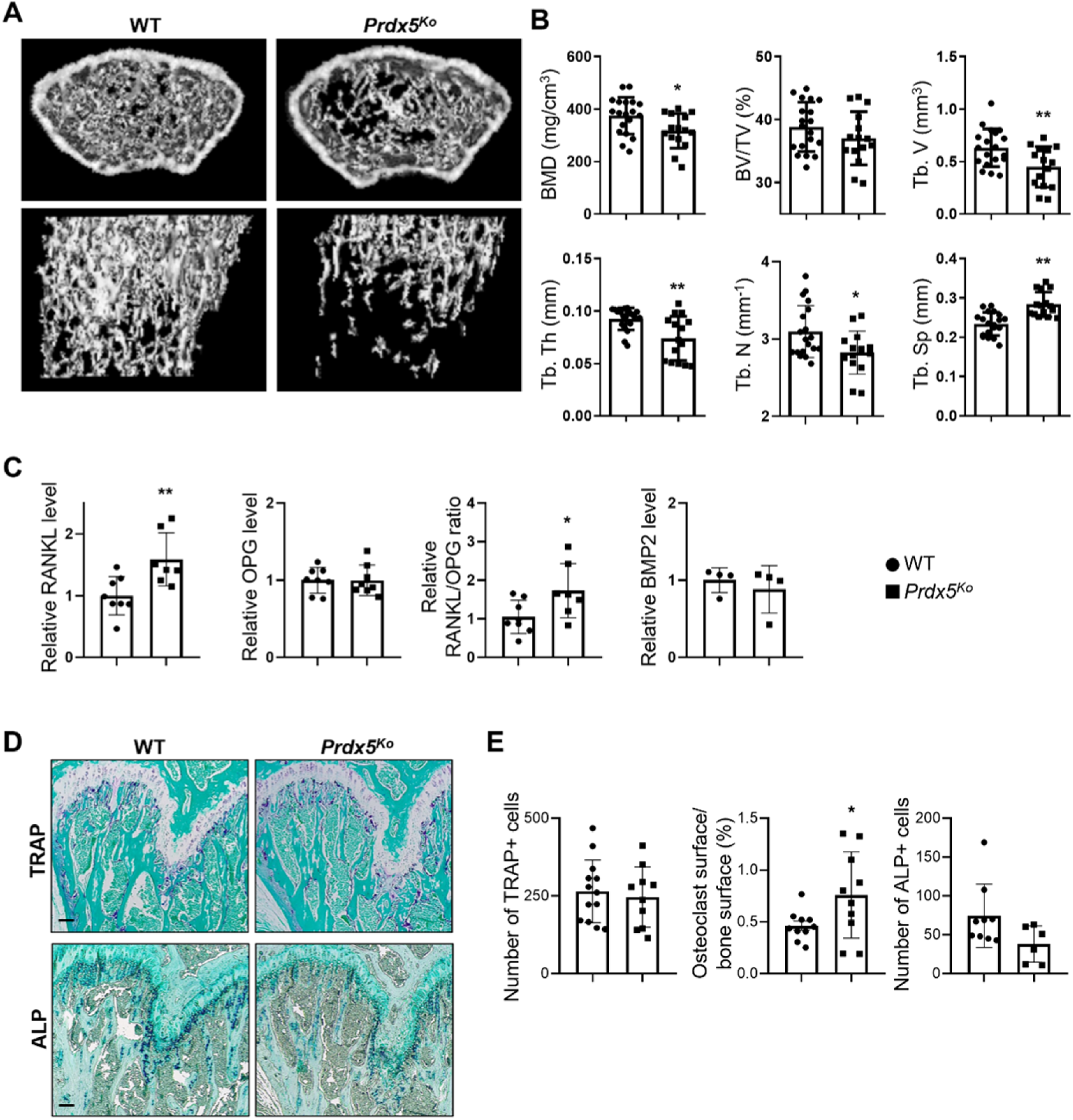
*Prdx5*^Ko^ mice show enhanced osteoporotic phenotypes. (A) Micro-CT images of femurs from 12-week-old WT and *Prdx5*^Ko^ male mice. (B) Micro-CT data were quantified (n = 15–19). BMD, bone mineral density; BV/TV, bone volume relative to total tissue volume; Tb. V, trabecular volume; Tb. Th, trabecular bone thickness; Tb. N, trabecular bone number; Tb. Sp, trabecular bone space. (C) Quantitative analysis of the levels of RANKL, OPG, and BMP2 in the sera from WT and *Prdx5*^Ko^ mice at 12 weeks. (n = 4–8). (D) Representative TRAP and ALP staining images of the mouse femora. TRAP- or ALP-positive cells were stained as purple, and the bone was counterstained with Fast Green as blue. Scale bar, 100 µm. (E) Quantification of the TRAP- or ALP-positive cells shown in (D). (n = 6–10)

We confirmed the osteogenic potential in mouse femurs stained with TRAP and ALP, which are the markers of osteoclasts and osteoblasts, respectively (Figure 3D). The number of total TRAP-positive cells was not altered in *Prdx5*^Ko^ mice. Since *Prdx5*^Ko^ mice showed less trabecular bone volume (Figure 3B), we measured the ratio of osteoclast and bone surfaces. *Prdx5*^Ko^ mice showed higher osteoclast surface ratios than WT (Figure 3E). The total number of ALP-positive cells reduced in *Prdx5*^Ko^ mice; however, the reduction was not statistically significant. Altogether, *Prdx5*^Ko^ mice showed increased number of osteoclasts in the femurs. These osteoporotic phenotypes were not observed in female mice (Figure 3– figure supplement 1). *Prdx5*^Ko^ females showed no differences in BMD, bone volume, and trabecular bone thickness and space. Therefore, we examined bone parameters in an ovariectomy-induced osteoporosis mouse model (OVX) (Figure 3–figure supplement 1). Micro-CT analysis revealed that OVX mice displayed significantly lower Tb. V and Tb. N than sham mice; however, no significant differences were observed between WT and *Prdx5*^Ko^ mice. These findings indicate that Prdx5 may not act as a hormone-dependent regulator, and osteoporosis phenotypes observed in males are specific responses of Prdx5. Nonetheless, hormones compensate for Prdx5-mediated osteoporosis.

### Limited bone remodeling activities in *Prdx5*^Ko^ mice

To determine the bone remodeling activity, we examined the bone turnover rates in *Prdx5*^Ko^ mice. First, we confirmed osteoblast function using trichrome staining and dynamic bone histomorphometry analysis *in vivo* (Figure 4–figure supplement 1). Trichrome staining revealed lower osteoid volume per bone volume in *Prdx5*^Ko^ mice than in WT, indicating reduced bone modeling in *Prdx5*-deficient mice. In *Prdx5*^Ko^ mice, a lower width between calcein and alizarin red S labeling and lower mineral apposition rate (MAR) in the trabecular bone were observed than those in WT mice. However, cortical bone revealed no alteration. Thus, *Prdx5*^Ko^ mice exhibited reduced bone turnover parameters, which indicated the suppression of newly formed bone tissue in the trabecular bone.

Next, to test osteogenic potential *in vivo*, we analyzed the osteogenic healing capacity using the calvarial defect model in *Prdx5*^Ko^ mice and their WT littermates (Figure 4). After the calvarial bone was trepanned, mice were treated with BMP2 or phosphate-buffered saline (PBS) for 3 weeks. In BMP2-administered mice, newly formed bones were observed; however, *Prdx5*^Ko^ mice showed a lesser extent of new bone formation than WT (Figure 4A). Immunostaining analysis was performed to measure cross-sectional area and bone volume. Larger puncture and smaller bone volume were observed in the calvaria of *Prdx5*^Ko^ mice than in those of WT (Figure 4B). The BMP2-restored lesions in *Prdx5*^Ko^ mice were thinner than those in WT mice, and the number of TRAP-positive cells and the osteoclast surface/bone surface ratio were similar in *Prdx5*^Ko^ and WT mice. However, *Prdx5*^Ko^ mice had fewer ALP-positive osteoblasts than WT (Figure 4C, D). These results imply that *Prdx5* plays an essential role in osteoblast-mediated bone regeneration.

**Figure 4.**
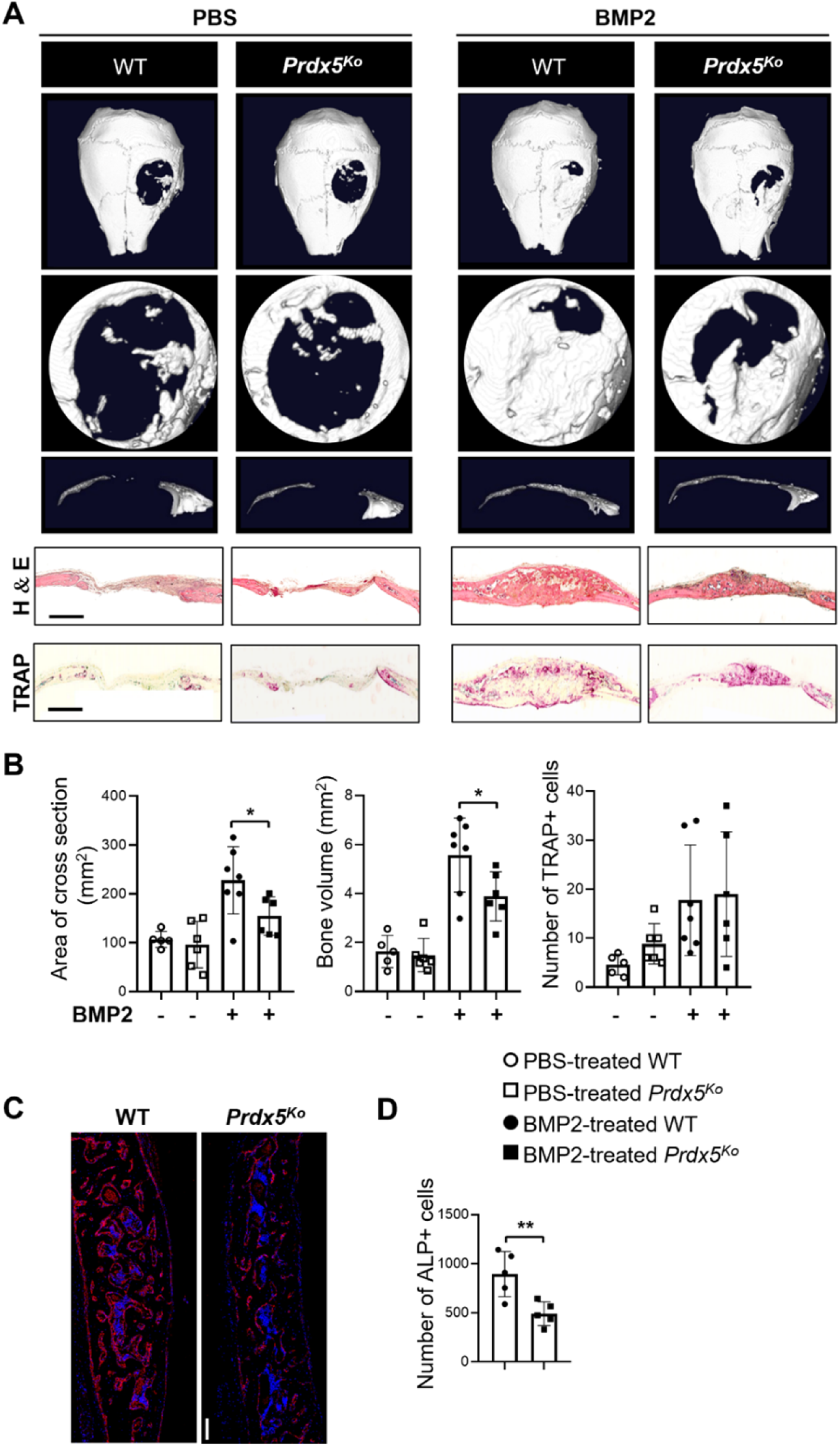
*Prdx5*^Ko^ mice show reduced bone healing after BMP2 induction. (A) Representative micro-CT images of the calvarial defect model after 3 weeks of implantation with PBS- or BMP2-containing sponges. The representative images show various shapes: whole (top), the hole from each image (middle); and the cross-section (bottom) from each hole. Representative hematoxylin–eosin and TRAP staining images of the calvarial bone section from each group. Scale bar, 1000 μM. (B) Measurement of the cross-sectional area, new bone formation, and number of TRAP-positive cells at the calvarial defect site (n = 5**–**7). (C) Representative images of ALP staining (scale bar, 100 μM) and (D) quantification of the number of ALP-positive cells (n = 5). ALP-positive cells were stained red, while DAPI-positive cells were counterstained blue.

### Prdx5 co-localizes and interacts with hnRNPK in response to BMP2 stimulation

Prdx5 expression increased during osteoblast differentiation (Figure 2), which suggested that Prdx5 acts as a positive regulator of osteoblast differentiation. To understand the role of Prdx5 in osteoblasts, we investigated Prdx5-interacting proteins using LC-MS/MS after immunoprecipitation with a Prdx5 antibody using *in vitro* differentiated osteoblasts (Figure 5A). We identified a total of 43 Prdx5-associated proteins (Table 1). In Gene Ontology (GO) analysis with these 43 proteins, RNA splicing was found to be the only significant biological pathway, suggesting the involvement of Prdx5 through an RNA-related mechanism (Figure 5B). To determine the interacting proteins responsive to BMP2, we focused on BMP2-specific proteins. A total of 20 proteins were classified as BMP2-specific interacting proteins (Figure 5A and Table 2). Since Prdx5 was localized in the nucleus after BMP2 stimulation (Figure 5–figure supplement 1), and to understand the function of Prdx5 in cell differentiation, we focused on nuclear proteins, hnRNPs. hnRNPK was found to be close to Prdx5 via STRING analysis (Figure 5C); this protein has previously been studied in osteoclasts and osteoblasts (Fan et al., 2015; Stains et al., 2005). Here, we confirmed the localization of Prdx5 and hnRNPK at the single-cell level (Figure 6A). After BMP2 stimulation, Prdx5 and hnRNPK were co-localized in the nucleus and cytosol in osteoblasts. We also confirmed the interaction between Prdx5 and hnRNPK using immunoprecipitation (Figure 6B). To clarify the relationship between Prdx5 and hnRNPK, we compared hnRNPK localization in *Prdx5*^Ko^ cells after BMP2 stimulation. Interestingly, hnRNPK was localized only in the nucleus in *Prdx5*^Ko^ cells, whereas it was observed in the cytosol and nucleus in WT (Figure 6C). To verify the microscopic data, the levels of hnRNPK were examined in the nuclear and cytoplasmic fractions of *Prdx5*^Ko^ osteoblasts (Figure 6D). Higher levels of hnRNPK were detected in the nuclear fraction of *Prdx5*^Ko^ osteoblasts than in that of the WT osteoblasts; the expression was similar in the absence of BMP2. These data suggest that Prdx5 may control the localization of hnRNPK in osteoblasts.

**Figure 5.**
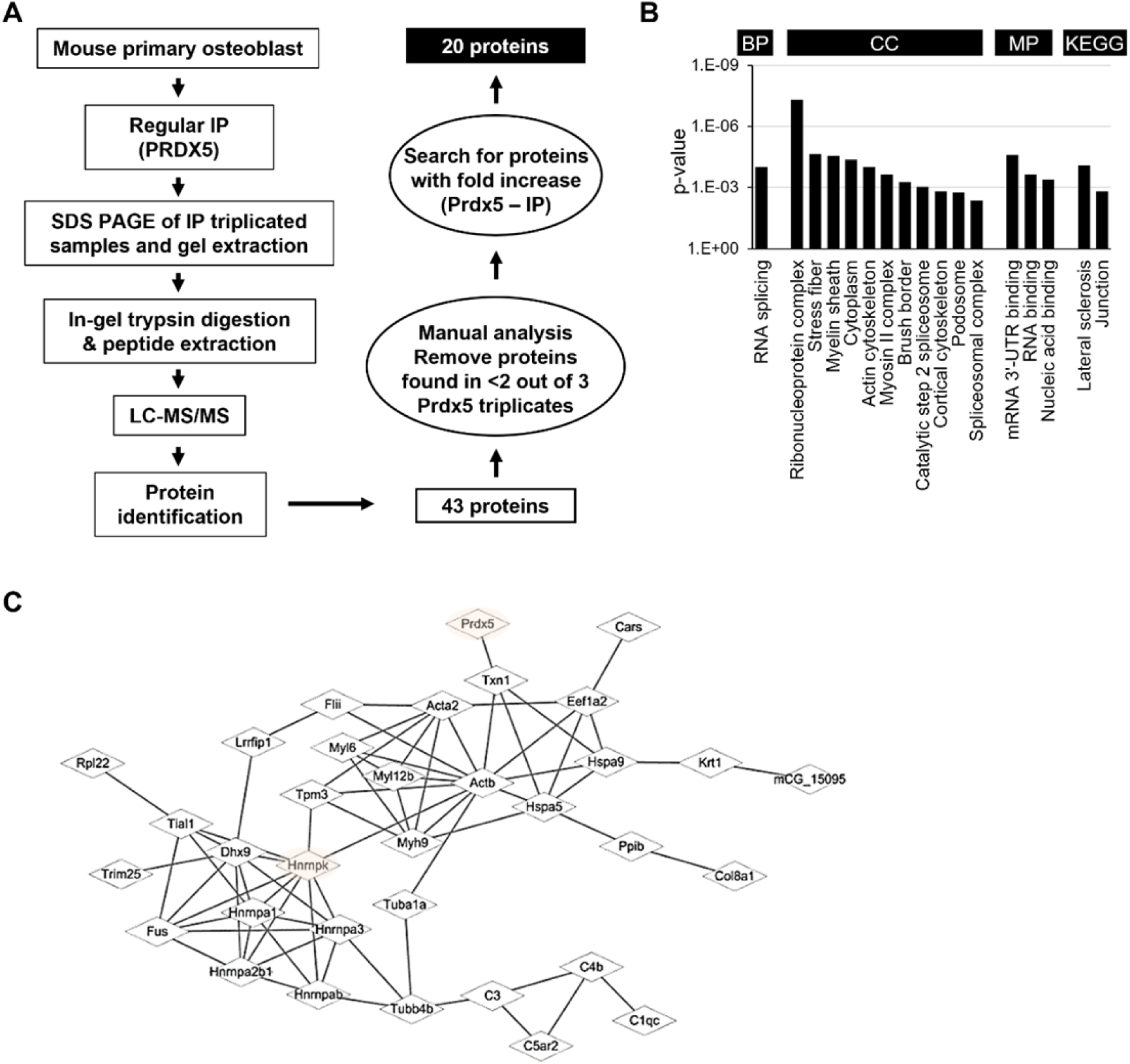
Identification of Prdx5-interacting proteins during osteoblast differentiation. (A) Schematic representation of the experimental design of IP and LC-MS/MS. Total 20 proteins were identified as potential candidates binding to Prdx5 in osteoclasts. (B) GO analysis results with 43 proteins are shown by biological process (BP), cellular component (CC), molecular function (MF), and Kyoto encyclopedia of genes and genomes (KEGG). (C) The interaction of Prdx5 with the 43 proteins identified in the MS/MS analysis was constructed using the STRING database.

**Figure 6.**
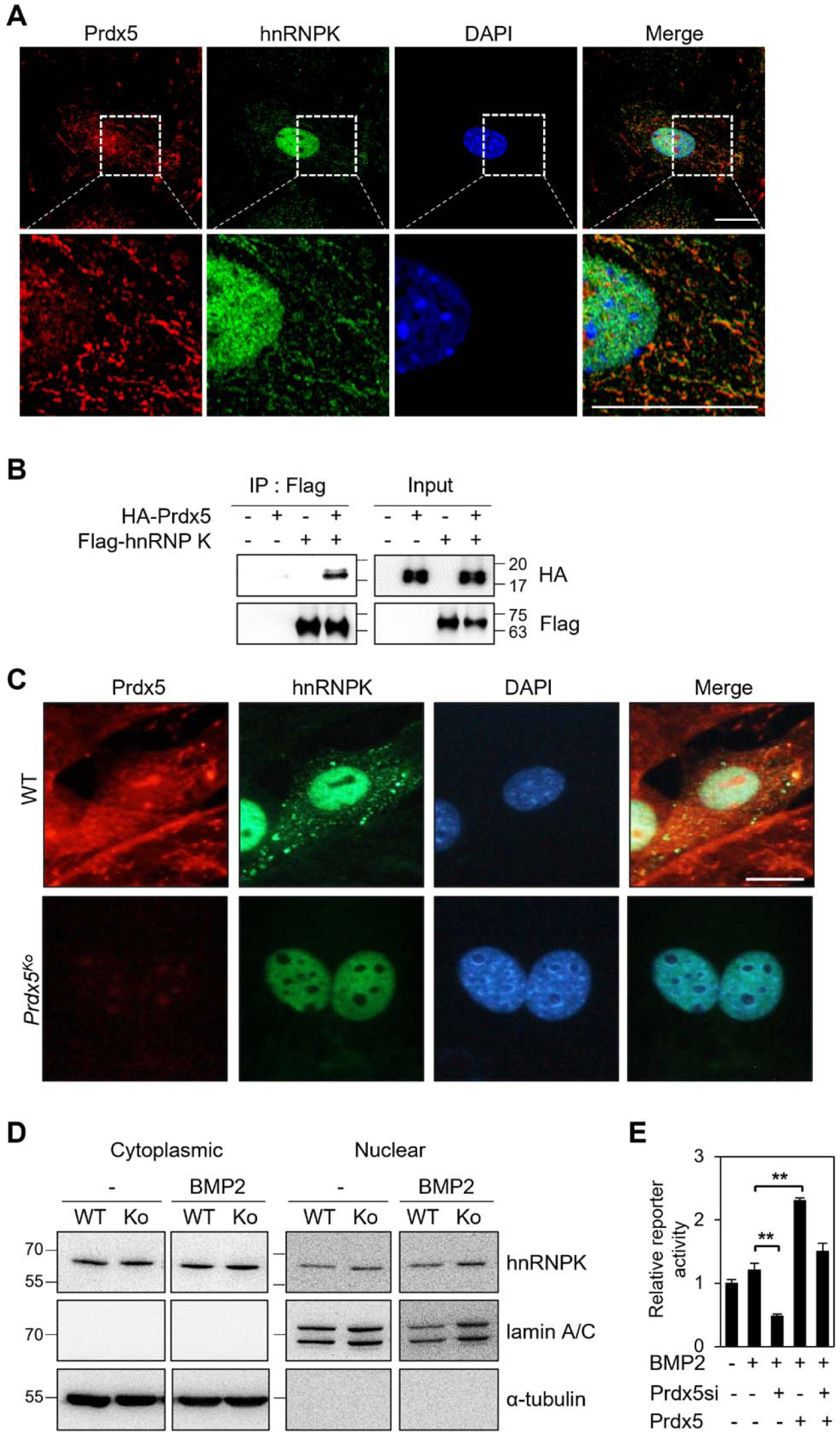
hnRNPK interacts with Prdx5 in osteoblasts. (A) To determine co-localization, osteoblasts were stained with antibodies against Prdx5 and hnRNPK, and images were acquired via confocal microscopy (scale bar, 20 μm). The upper images were magnified as depicted by the dotted box in the lower images. (B) Immunoprecipitation was performed using HEK293T cells expressing various combinations of HA-tagged Prdx5 and flag-tagged hnRNPK. (C) Osteoblasts were differentiated from precursors derived from WT and *Prdx5*^Ko^ mice via BMP2 treatment for 7 days. hnRNPK localization was analyzed via confocal microscopy (scale bar, 20 µm). (D) hnRNPK levels were determined in the cytoplasmic and nuclear fractions of WT and *Prdx5*^Ko^ cells. Osteoblasts were harvested on day 7. (E) The OG2-luciferase assay was performed using MC3T3-E1 cells differentially expressing Prdx5 and BMP2 stimulation.

**Table 1.**
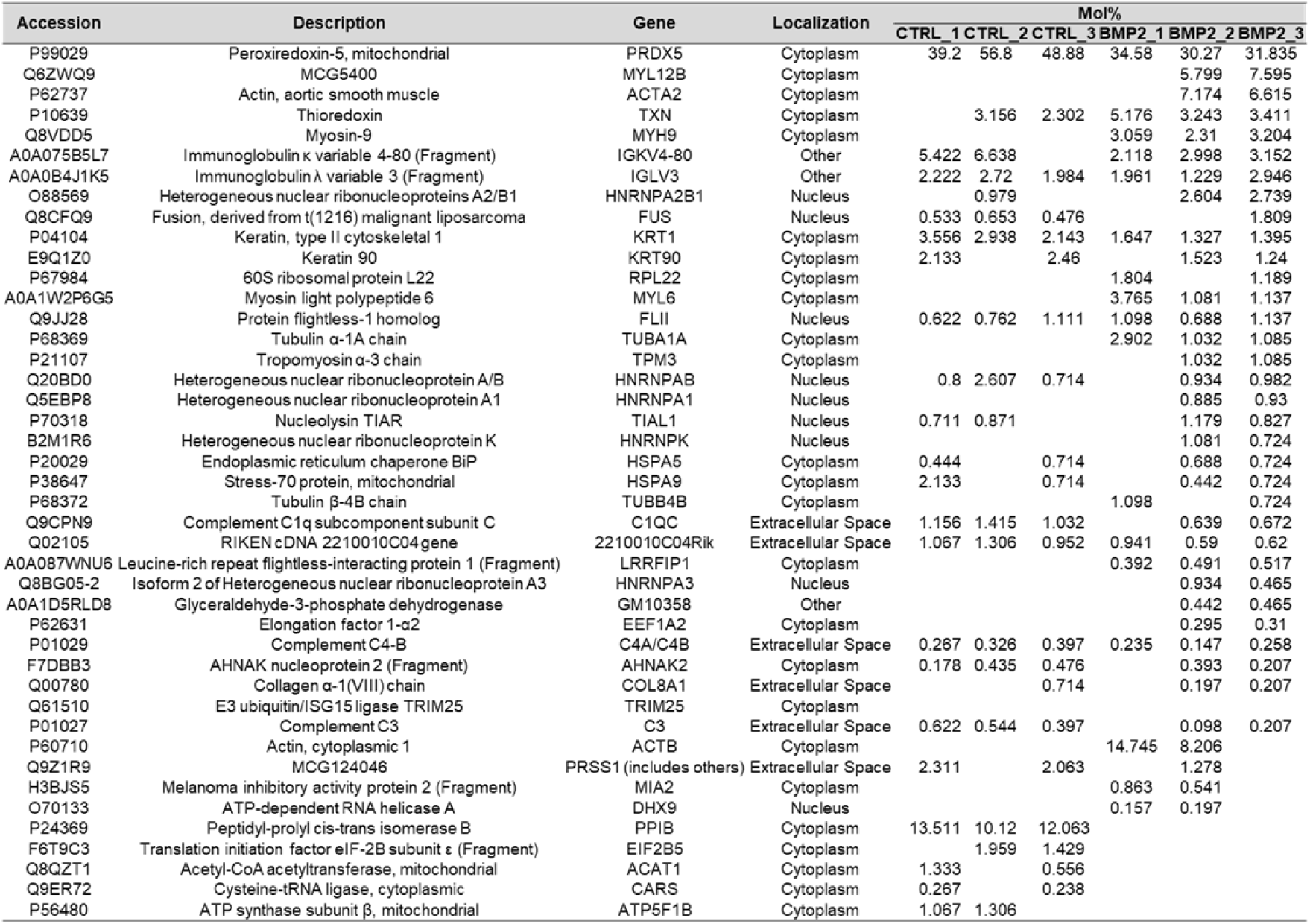
The 43 Prdx5-interacting proteins identified via LC-MS/MS analysis

**Table 2.**
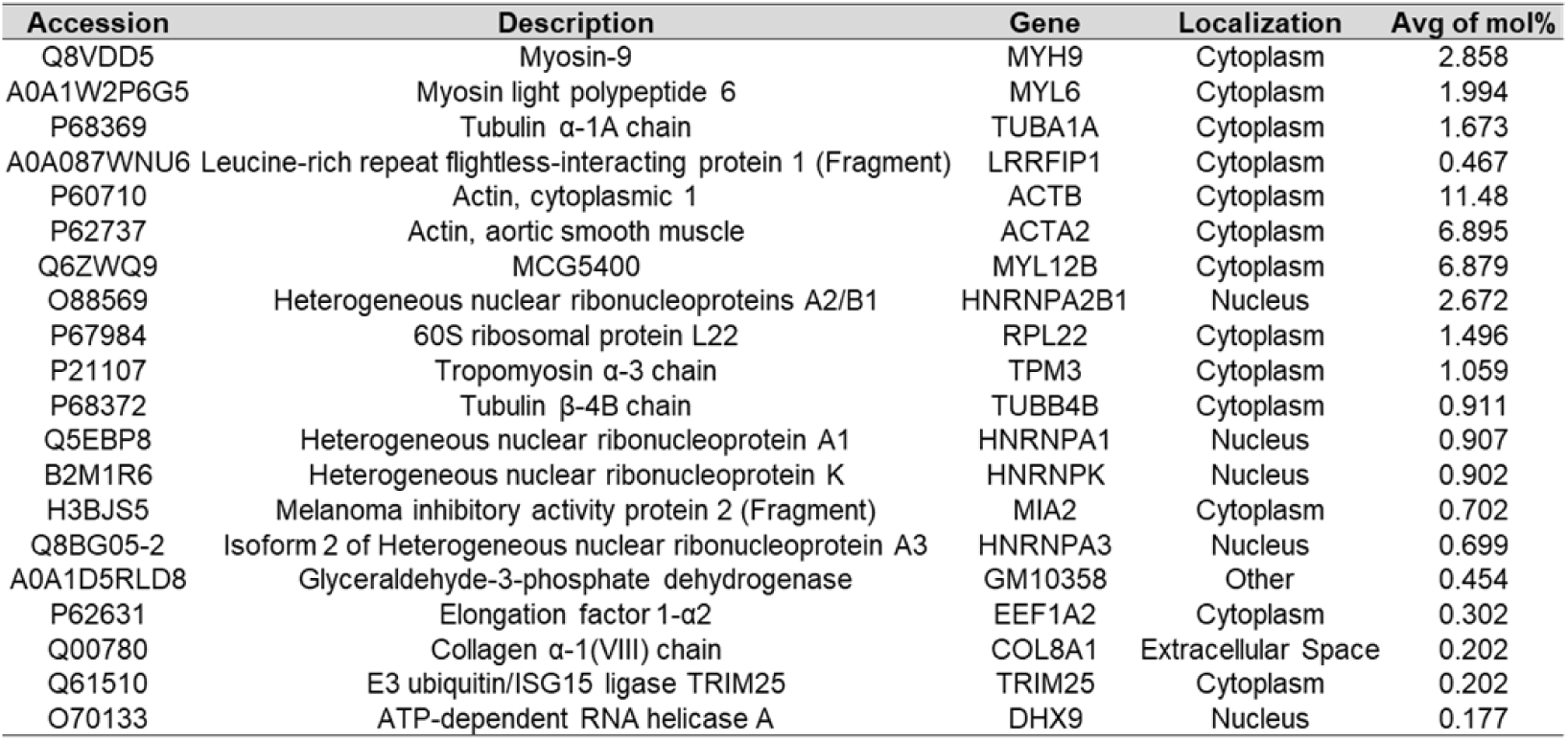
Prdx5-interacting proteins detected only in the BMP2-treated group

In osteoblasts, previous studies have reported the role of hnRNPK as a repressor of *Ocn* expression (Niger et al., 2011; Stains et al., 2005). Since Prdx5 acted as an activator of osteoblast differentiation in our study, and *Ocn* levels were attenuated in osteoblasts from *Prdx5*^Ko^ mice (Figure 2C), we assumed that Prdx5 inhibits hnRNPK to regulate *Ocn*. We performed a reporter assay using *Ocn* promoter to verify whether Prdx5 affects *Ocn* expression (Figure 6E). We found that Prdx5 knockdown suppressed the Ocn activity that was rescued by Prdx5 overexpression. Altogether, these results indicate that Prdx5 interacts with hnRNPK in osteoblasts to transport hnRNPK from the nucleus to cytoplasm, which activates *Ocn* to induce osteoblast differentiation.

### Expression of osteoclast-related genes is increased in *Prdx5*^Ko^ osteoclasts

As *Prdx5*^Ko^ mice showed an increase in the number of TRAP-positive osteoclasts in the femurs (Figure 3E), and BMMs from *Prdx5*^Ko^ mice differentiated into more osteoclasts than those in WT (Figure 2E), we analyzed the transcriptome profiles of BMMs and osteoclasts via RNA-seq (Figure 7). BMMs were isolated from WT and *Prdx5*^Ko^ mice, and the cells were differentiated into osteoclasts via RANKL stimulation for 4 days *in vitro*. The number of reads raged from 72,748,470 to 86,717,526 was generated, and the trimmed clean reads were mapped to the mouse reference genome with 97–98 % alignment rates (Figure 7–table supplement 1). BMMs and osteoclasts were clearly separated by principal component analysis (PCA) (Figure 7A). However, no significant differences were observed between the BMMs of WT and *Prdx5*^Ko^ mice. A comparison of differentially expressed genes (DEGs) between WT and *Prdx5*^Ko^ cells revealed 214 DEGs in BMMs, whereas 1257 genes were detected in osteoclasts (Figure 7B, C). Among the 214 genes, 61 (28.5%) were upregulated and 153 (71.5%) were downregulated in *Prdx5*^Ko^ mice compared to those in WT. However, approximately half of DEGs were up- and downregulated in *Prdx5*^Ko^ osteoclasts (51% and 49%, respectively). These results suggest that Prdx5 acts as an activator of gene expression in BMMs, and the gene levels are high in osteoclast precursors. However, the levels of these genes decrease during osteoclastogenesis. In GO analysis, the DEGs were found to be involved in the immune response (Figure 7–figure supplement 1).

**Figure 7.**
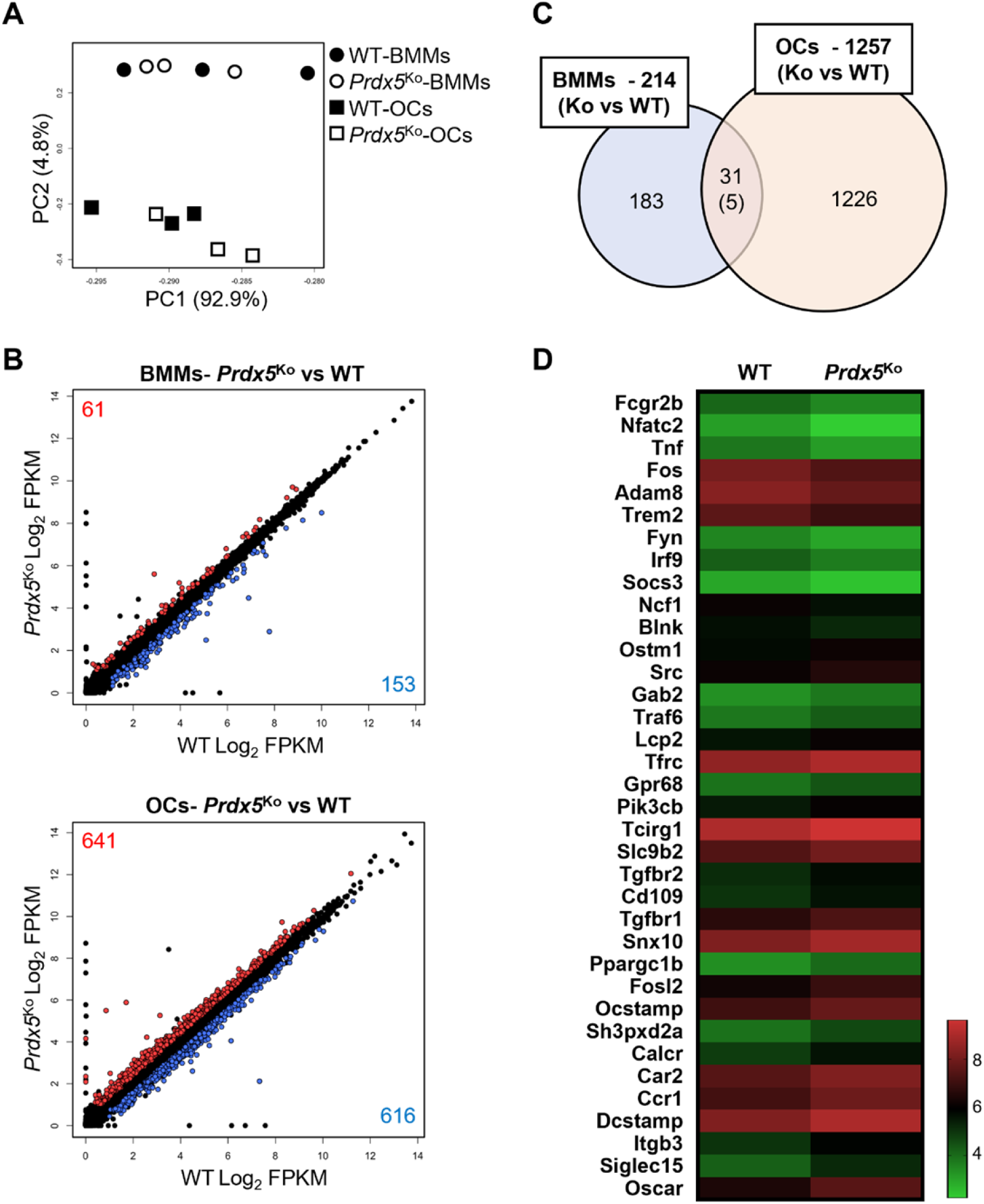
Osteoclast-related genes are highly expressed in *Prdx5*-deficient osteoclasts. (A) PCA of BMMs and osteoclasts (OCs) from WT and *Prdx5*^Ko^ cells. Each circle or square represents the expression profile of one sample (n = 3). (B) The DEGs in BMMs and OCs by comparison of *Prdx5*^Ko^ versus WT are displayed on a scatter plot. Each dot indicates a single gene. Significantly upregulated DEGs in *Prdx5*^Ko^ are indicated in red, while downregulated DEGs are indicated in blue (FPKM > 1, q-value < 0.05). (C) The Venn diagram indicates DEGs in BMMs and OCs. A total of 31 DEGs are overlapped in BMMs and OCs, and only five genes show opposite patterns, which are downregulated in *Prdx5*^Ko^ OCs but upregulated in *Prdx5*^Ko^ BMMs. (D) Heatmap analysis shows the osteoclast-related DEGs. The z-score represents log_2_ FPKM.

We hypothesized that *Prdx5* deficiency results in a positive regulation of osteoclast differentiation. In GO analysis, the downregulated DEGs in *Prdx5*^Ko^ osteoclasts were involved in cell cycle regulation and cell division, while the upregulated DEGs were enriched in signaling and osteoclast differentiation (Figure 7–figure supplement 1). When we examined osteoclast-related genes, 25 out of 36 DEGs were upregulated in *Prdx5*^Ko^ osteoclasts (Figure 7D). Interestingly, the levels of transcription factors (*NFATc1*, *Fos*, and *Irf9*) that regulated the early response of osteoclast differentiation were suppressed in *Prdx5*^Ko^ osteoclasts. In contrast, osteoclast maker genes (*OC-STAMP*, *Calcr*, *DC-Stamp*, *Itgb3*, and *Oscar*), which are highly expressed in mature osteoclasts, were upregulated in *Prdx5*^Ko^ osteoclasts.

## Discussion

Osteoporosis results in an excessive reduction in bone mass, which is a major health issue in the elderly population (Demontiero et al., 2012). Clinically, some therapeutic treatments are available to induce osteoblast and reduce osteoclast activities (Milat & Ebeling, 2016). However, these treatments are associated with severe side effects, including heart issues, kidney damage, and osteonecrosis of the jaw (Compston et al., 2019; Saag et al., 2017). Therefore, a novel drug with curative and fewer side effects is urgently needed to treat osteoporosis.

Here, we assessed the critical functions of Prdx5 in bone homeostasis. Prdx5 expression increased during osteoblast differentiation and decreased during osteoclast differentiation. Genetically deficient *Prdx5* mice developed osteoporosis-like phenotypes, which suggests that Prdx5 is important in bone remodeling. In osteoblasts, Prdx5 and hnRNPK were co-localized in the nucleus and cytosol, and Prdx5 regulated the hnRNPK-mediated *Ocn* transcription. In osteoclasts, Prdx5 acted as an inhibitor, as revealed by the upregulation of osteoclast-related genes in *Prdx5*^ko^ cells. We demonstrated that Prdx5 was a novel positive regulator of osteoblast differentiation, and that it also regulated osteoclastogenesis. Our study indicated the beneficial pharmacological effect of Prdx5 in the maintenance of bone mass during the formation of skeletal tissues.

Six members of the Prdx family reportedly exhibit antioxidant activities owing to the presence of CXXC amino acid sequences (Chae et al., 1994; Rhee et al., 2001). Prdx5 is a unique member of the atypical 2-Cys subfamily in mammals and is expressed ubiquitously in all tissues (Rhee et al., 2001). Prdx5 is present in the cytosol, peroxisomes, and mitochondria (Rhee et al., 2012). *Prdx5* deficiency leads to an increased susceptibility to high-fat diet-induced obesity and metabolic abnormalities (Kim et al., 2018; Kim et al., 2020). In this study, we first investigated the changes in osteogenesis or bone mass formation by Prdx5. In addition, we confirmed the role of Prdx5 in osteogenic processes. The *Prdx5*^Ko^ mice showed a significant reduction in bone mass, which suggested that Prdx5 affected bone turnover. *Prdx5* deficiency markedly inhibited osteoblast differentiation and increased osteoclast differentiation *in vitro*. Indeed, the bone healing rate and osteocyte population decreased in *Prdx5*^Ko^ mice. Interestingly, Prdx5 may interact with hnRNPK in osteoblasts. Given the reduced bone mass in *Prdx5*^Ko^ mice, we investigated the function of Prdx5 in osteoclasts. However, we did not focus on the role of Prdx5 in osteoclasts, because its expression was extremely low after RANKL stimulation. Our results imply that Prdx5 primarily acts in the osteoblasts, and it may not be necessary for osteoclasts.

To determine the antioxidative role of Prdx5 in bone cell differentiation, we determined the ROS levels in BMP2-treated osteoblasts. ROS production was not altered by BMP2 stimulation in *Prdx5*-deficient cells. We found that Prdx5 is involved in ROS generation during osteoblast differentiation, which is necessary for BMP2-mediated ROS production. However, this mechanism is an early response in the cytoplasm, which is scavenged later during osteoblastogenesis. Further studies are required to elucidate the relationship between ROS and Prdx5 in bone cells, particularly, in terms of mitochondrial functions. In this study, we primarily focused on the role of Prdx5 in the nucleus.

hnRNPK also interacts with numerous proteins in the nucleus and cytosol, including signal transduction proteins, transcriptional activators, and repressors (Naarmann et al., 2008; Perrotti & Neviani, 2007; Ritchie et al., 2003). Therefore, hnRNPK may act as a docking platform or scaffold, shuttling from the cytoplasm to the nucleus (Krecic & Swanson, 1999; Mikula et al., 2010). Prdx5 was also expressed in the cytosol and nucleus (Figure 7). Furthermore, we also examined Prdx5 translocation to the nucleus upon BMP2 induction. Our results suggested potential mechanisms through which transcriptional repression by hnRNPK may occur. The most likely scenario is that hnRNPK competitively binds to an unknown transcription factor (complex II) that binds to the putative CT-rich region of the *Ocn* promoter, resulting in the loss of an activator from the promoter and a net repression of gene transcription (Stains et al., 2005).Our results indicated that Prdx5 disturbed the binding potential of hnRNPK to suppress *Ocn* expression through an interaction between Prdx5 and hnRNPK and their translocation. hnRNPK interacts with glycogen synthase kinase-3b during osteoclast differentiation via nuclear–cytoplasmic translocation (Fan et al., 2015). Further studies may demonstrate the correlation between Prdx5 attenuation and hnRNPK translocation during osteoclastogenesis.

In conclusion, we identified a new mechanism of Prdx5 in regulating the hnRNPK–Ocn axis in osteoblasts. Our study also indicates that Prdx5 controls osteoclast differentiation, which is mediated by osteoblast differentiation or the early stages of osteoclastogenesis. Therefore, Prdx5 is critical in bone remodeling.

## Materials and Methods

### Key resources table

| Reagent type (species) or resource | Designation | Source or reference | Identifiers | Additional information |
| --- | --- | --- | --- | --- |
| Genetic Reagent (M. musculus) | C57BL/6 J | Kim *et al.*, 2018 |  | DBL Co. Daejeon, South Korea |
| Chemical compound | BMP2 | Sino Biological | Cat#10426 |  |
| Chemical compound | Recombinant murine sRANKL | PeproTech | Cat#315-11 |  |
| Chemical compound | Recombinant murine M-CSF | PeproTech | Cat#315-02 |  |
| Chemical compound | Type I collagenase | Gibco | Cat#17018 |  |
| Chemical compound | Dispase | Roche | Cat#4942078001 |  |
| Chemical compound | BCIP/NBT alkaline phosphatase kit | Sigma-Aldrich | Cat#B1911 |  |
| Chemical compound | CM-H2DCFDA | Thermo Fisher Scientific | Cat#MP36103 |  |
| Chemical compound | Calcein | Sigma-Aldrich | Cat#C0875 |  |
| Chemical compound | Alizarin S Red | Sigma-Aldrich | Cat#A5533 |  |
| Chemical compound | Prdx5 siRNA | Thermo Fisher Scientific | Cat#s79839-41 |  |
| Chemical compound | Lipofectamine 3000 | Invitrogen | Cat#L3000001 |  |
| Chemical compound | Hematoxylin | Sigma-Aldrich | Cat#H9627 |  |
| Chemical compound | Bouin's solution | Sigma-Aldrich | Cat#HT10132 |  |
| Chemical compound | Acid Fuchsin | Sigma-Aldrich | Cat#F-8129 |  |
| Chemical compound | Aniline blue | Junsei | Cat#11466 |  |
| Chemical compound | Phosphomolybdic acid hydrate | Junsei | Cat#84235 |  |
| Chemical compound | Phosphotungstic acid hydrate | Junsei | Cat#84220 |  |
| Chemical compound | Fast Green FCF | Sigma-Aldrich | Cat#7252 |  |
| Commercial assay or kit | TRAP Staining Kit | Cosmo Bio Co. | Cat#PMC-AK04-COS |  |
| Commercial assay or kit | SYBR Green Master mix | Thermo Fisher Scientific | Cat#A25778 |  |
| Commercial assay or kit | Reverse transcription kit | Thermo Fisher Scientific | Cat#18064022 |  |
| Commercial assay or kit | Mouse RANKL ELISA | Abcam | Cat#ag100749 |  |
| Commercial assay or kit | Mouse OPG ELISA | R&D systems | Cat#MOP00 |  |
| Commercial assay or kit | Mouse BMP2 ELISA | LS bio | Cat#LS-F36595 |  |
| Commercial assay or kit | Bone Resorption assay kit | Cosmo Bio Co. | Cat#CSR-BRA |  |
| Commercial assay or kit | Nuclear and cytoplasmic extraction kit | Thermo Fisher Scientific | Cat#78833 |  |
| Commercial assay or kit | Luciferase assay system | Promega | Cat#E1500 |  |
| Antibody | Mouse monoclonal anti-Prdx5 | Invitrogen | Cat#LF-MA0002 | (1:1000) |
| Antibody | Rabbit polyclonal anti-Prdx5 | Ab Frontier | Cat#LF-PA0010 | (1:500) |
| Antibody | Rabbit polyclonal anti-hnRNPK | Cell Signaling Technology | Cat#9081 | (1:1000) |
| Antibody | Mouse monoclonal anti-beta actin | Sigma | Cat#A5441 | (1:1000) |
| Antibody | Anti-ALP | Abcam | Cat#ab229126 | (1:200) |
| Antibody | HA-Tag | Santa Cruz | Cat#sc-7392 | (1:1000) |
| Antibody | Flag (OctA)-probe | Santa Cruz | Cat#sc-166355 | (1:1000) |
| Antibody | Lamin A/C | Cell Signaling Technology | Cat#2032 | (1:1000) |
| Antibody | Lamin B | Ab Frontier | Cat#LF-PA50043 | (1:1000) |
| Antibody | Tubulin alpha | Novus | Cat#NB110 | (1:1000) |
| Antibody | Anti rabbit Alexa Fluor 488 | Thermo Fisher Scientific | Cat#A-32731 | (1:200) |
| Antibody | Anti mouse Alexa Fluor 555 | Thermo Fisher Scientific | Cat#A-21427 | (1:200) |
| Software Algorithm | Graphpad 8 | https://www.graphpad.com/ |  |  |

### Animal experiments

All animals were housed in a specific pathogen-free facility following the guidelines provided in the Guide for the Care and Use of Laboratory Animals (Chonnam National University, Gwangju, Korea). All animal experiments were approved by the Institutional Animal Care and Use Committee (IACUC) of Chonnam National University (Approval No. CNU IACUC-YB-2019-50, CNU IACUC-YB-2017-53), Gwangju, Republic of Korea.

*Prdx5*^Ko^ (C57BL/6J) mice were gifted by Dr. Hyun-ae Woo, Ewha Womans University, Republic of Korea (Kim et al., 2018). To obtain the WT and transgenic mice, heterozygous males and females were crossed, and littermates were used for experiments.

Eight-week-old WT and their transgenic female littermates were sham-operated or subjected to bilateral OVX under anesthesia (25 mg/kg Zoletil and 12.5 mg/kg Rompun). The mice were sacrificed after 4 weeks, and serum, uterus, and femurs were collected for biochemical and histomorphometric analyses.

### Osteoclast and osteoblast differentiation *in vitro*

Primary mouse pre-osteoblasts were isolated from the calvaria of 3-day-old C57BL/6J mice via sequential digestion with type I collagenase (Gibco) and dispase (Roche), as previously described (Bellows et al., 1986). Briefly, the cells were cultured in an α-minimum essential medium (α-MEM), containing 10% characterized heat-inactivated fetal bovine serum (FBS) and 1% penicillin/streptomycin, and differentiated into osteoblasts via treatment with 100 ng/mL BMP-2 (Sino Biological). Cells were harvested at indicated time periods, and ALP staining was performed on day 7. For ALP staining, cells were fixed in 70% ethanol for 1 h and stained for 10 min with an ALP staining solution (BCIP/NBT alkaline phosphatase kit, Sigma-Aldrich), according to the manufacturer’s instructions.

For *in vitro* osteoclast differentiation, bone marrow-derived macrophage cells were isolated and stimulated with 30 ng/mL M-CSF (PeproTech) and 50 ng/mL RANKL (PeproTech) as previously described (Cho et al., 2021). To assess the extent of differentiation, the cells were stained using a TRAP kit (Cosmo Bio Co.). The mature osteoclasts were counted under a microscope based on the number of nuclei (n ≥3), cell size, and cell number.

### Western blot analysis and qRT-PCR

The differentiated osteoblasts and osteoclasts were lysed in a radioimmune assay precipitation buffer (Thermo Scientific), and western blotting was performed as described previously (Cho et al., 2021). Mouse anti-Prdx5 (Invitrogen), rabbit anti-hnRNPK (CST), rabbit anti-Lamin β (Ab Frontier), and mouse anti-β-Actin (Sigma-Aldrich) antibodies were used to detect proteins.

Total RNA was extracted using TRIzol reagent (Thermo Fisher Scientific), and cDNA was synthesized as previously described (Cho et al., 2021). Quantitative PCR was performed using a SYBR Green-based system (Thermo Fisher Scientific), and data were calculated using the 2^-ΔΔCT^ method. Three separate experiments were performed. The primers used are listed in Table 3.

**Table 3.**
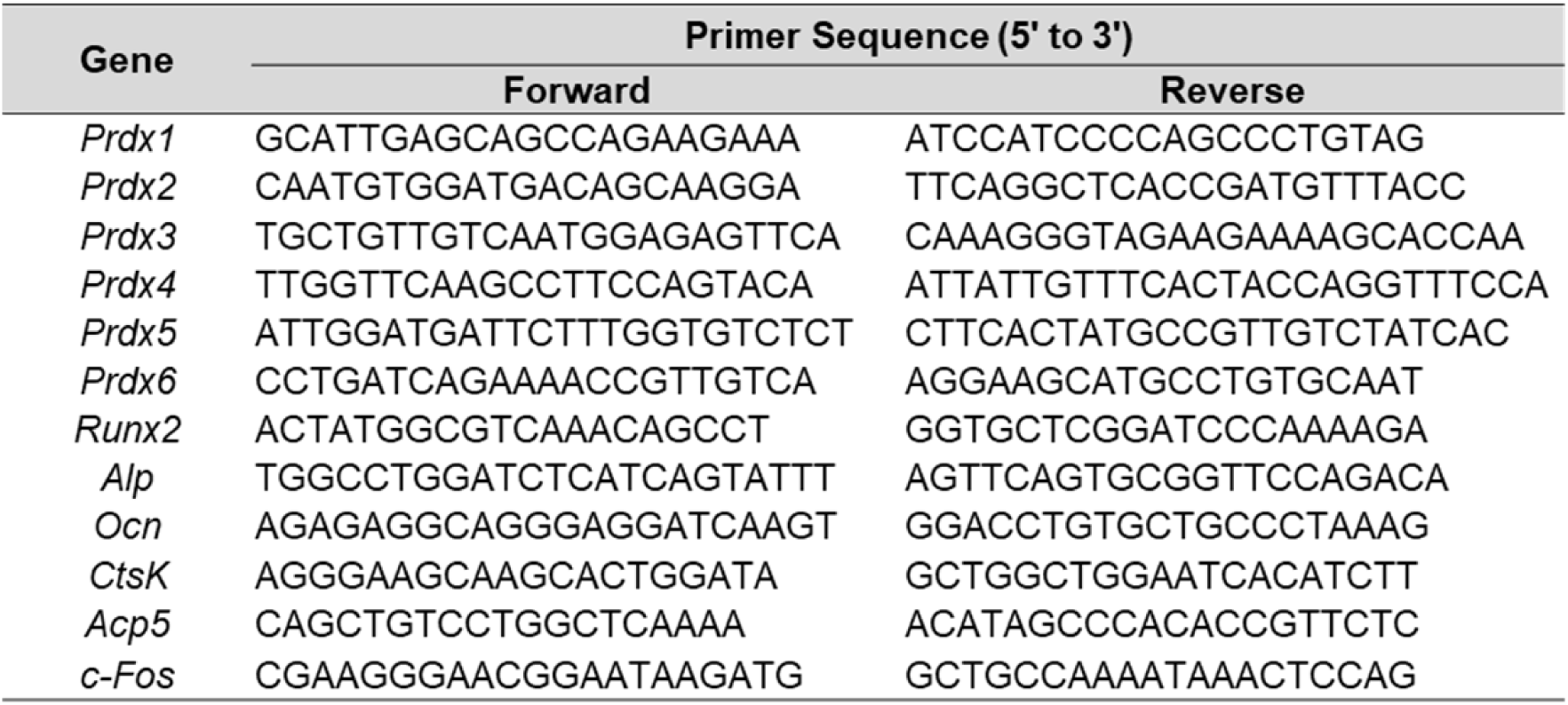
Primer sequences for qRT-PCR

### Micro-CT analysis

Femoral specimens were fixed in a 4% paraformaldehyde solution for 12 h at 4 °C, and micro-CT imaging was performed using a high-resolution Skyscan 1172 system (Bruker-micro-CT, Kontich, Belgium). The images were acquired at a 7 μm voxel resolution, with a 0.5 mm aluminum filter, at 50 kV and 100 μA exposure time, 0.5° rotation, and frame averaging of 1. An image reconstruction software (NRecon; Bruker) was used to reconstruct serial cross-section images using identical thresholds for all samples. For measuring the regions of interest (ROIs) of the trabecular and cortical bones, we included ROIs that were 0.7–2.3 mm away from the bottom of the growth plate. The bone morphometric parameters were calculated using adaptive thresholding (the mean of the minimum and maximum values) with CT Analyzer (version 1.11.8.0).

### Histology, immunostaining, and dynamic bone histomorphometry

Dynamic bone histomorphometric analysis was performed after injecting 25 mg/kg calcein or alizarin red (AR) into mice as previously described (Lim et al., 2015). Briefly, the distal femurs were fixed in a 4% paraformaldehyde solution and subsequently dehydrated with graded ethanol solutions; the undecalcified femurs were embedded in methyl methacrylate to prepare resin blocks. The resin blocks were cut longitudinally into 6 μm slices of the femur distal metaphysis using a Leica SP1600 microtome (Leica Microsystems, Germany). Fluorescence signals of calcein and AR from the ROIs were captured using a fluorescence microscope (Q500MC, Leica Microsystems). The parameters for dynamic bone histomorphometry were determined using the Bioquant Osteo 2018ME program (Bioquant Osteo, Nashville, TN, USA).

Goldner’s trichrome staining was performed on paraffin-embedded sections of 3 μm in length. After rehydration, the slides were washed in distilled water, refixed in Bouin’s solution (Sigma-Aldrich) for 15 min at 56 °C, and rinsed with running tap water for 5 min to remove picric acid (yellow color). The slides were counterstained with Weigert’s hematoxylin (Sigma-Aldrich) for 10 min, washed with tap water for 5 min, and rinsed thrice with distilled water. The slides were then stained with Biebrich scarlet-acid fuchsin (Sigma-Aldrich) for 5 min and rinsed thrice with distilled water. Next, the slides were immersed in phosphotungstic/phosphomolybdic acid (Junsei) for 10 min and transferred to aniline blue solution (Junsei) for 5 min. Finally, the slides were washed with distilled water and treated with 1% acetic acid for 1 min. After dehydration and mounting, the stained bone sections were observed under a microscope (Q500MC, Leica Microsystems), and the parameters of osteoid volume/bone volume were determined using the Bioquant Osteo 2018ME program (Bioquant Osteo).

Osteoclasts and osteoblasts were visualized using TRAP and ALP staining, respectively. TRAP (TRAP Staining Kit, Cosmo Bio Co.) staining was carried out according to the manufacturer’s instructions, with some modifications. NBT/BCIP staining (Sigma-Aldrich) was carried out by incubating tissue sections. The sections were then counterstained with 0.05 % Fast Green FCF (Sigma-Aldrich), dehydrated using graded ethanol solutions, and allowed to dry without clearing in xylene before mounting. Positive cells were visualized by purple color and analyzed using the ImageJ software.

### Enzyme-linked immunosorbent assay (ELISA)

The levels of specific markers of osteogenesis in the serum were measured using ELISA according to manufacturer’s description. RANKL levels were measured using a mouse RANKL ELISA kit (Abcam); OPG levels were measured using a Quantikine ELISA (R&D system) kit; BMP2 levels were measured using mouse BMP2 ELISA kits, respectively (LS Bio).

### Measurement of intracellular ROS levels

For osteoblasts, calvarial cells from WT and *Prdx5*^Ko^ mice were cultured for two days in a medium containing BMP2. For osteoclasts, BMMs from WT and *Prdx5*^Ko^ mice were cultured for two days in a medium containing M-CSF and RANKL. The cells were washed with α-MEM lacking phenol red and then incubated with 10 µM CM-H_2_DCFDA (Thermo Fisher Scientific) for 30 min. Fluorescence intensity was measured using a multiplate reader (SpectraMax i3x, Molecular Devices) and visualized under a microscope (Olympus Corp., IX2-ILL100) at excitation and emission wavelengths of 490 and 520 nm, respectively.

### Calvarial bone defect models and micro-CT analysis

For the calvarial bone defect model, a critical size calvarial defect was created using a 5 mm diameter trephine bur (Fine Science Tools, Foster City, CA, USA) and covered with absorbable collagen sponges containing 300 ng BMP-2 (Cowell Medi Corp., Seoul, Republic of Korea) in 12-week-old *Prdx5*^Ko^ and WT C57BL6/J male mice. After three weeks, the model mice were sacrificed for analysis. Briefly, the mice were subjected to inhalational anesthesia using an XGI-8 Gas Anesthesia System (PerkinElmer, Waltham, MA, USA) containing a mixture of 4% isoflurane (ISOTROY 100, Troikaa, India) and oxygen, for 4 min. The osteological structures of the specimens were examined using a micro-CT scanning system, combined with a Quantum GX µCT imaging system (PerkinElmer), at the Korea Basic Science Institute (Gwangju, Republic of Korea). The scanned skeletal data were reconstructed into 3D tomograms comprising high-contrast images of the skeletal parts of interest.

### Confocal microscopy

The cells were grown on sterilized glass coverslips and fixed in 4% paraformaldehyde. Non-specific binding was blocked by incubation of slides in 0.1% bovine serum albumin in PBS. Subsequently, the samples were stained with mouse anti-Prdx5 (1:200, Invitrogen) and rabbit anti-hnRNPK antibodies (1:200, Cell Signaling Technology), followed by incubation with Alexa 555- or Alexa 488-conjugated secondary antibodies (1:500, Invitrogen) and DAPI/antifade (1:200, Invitrogen). Images were captured using a confocal laser scanning microscope equipped with visible and near-infrared lasers. Images were acquired using the Airyscan mode supported by the LSM 880 confocal laser scanning microscope for image optimization (Carl Zeiss, Oberkochen, Germany).

### Immunoprecipitation (IP)

Pre-osteoblasts isolated from mouse calvaria were cultured for 7 days in a BMP2-containing or normal medium (CTRL). The cells were lysed with an IP lysis buffer (150 mM NaCl, 25 mM Tris-HCl, 10% glycerol, and 1 mM EDTA) containing a protease inhibitor cocktail (Roche, Basel, Switzerland). IP was performed with an anti-Prdx5 antibody (Ab Frontier). Sodium dodecyl sulfate-polyacrylamide gel electrophoresis and in-gel digestion were performed as previously described (Yun et al., 2018). Briefly, the sliced gel was dried and digested in trypsin. The tryptic peptides were dried and extracted for LC-MS/MS.

The lysed cells were centrifuged, and equal amounts of proteins were incubated with an anti-Prdx5 antibody, or an IgG rabbit polyclonal antibody (Cell Signaling Technology) as a negative control. The proteins were further incubated with protein A/G-sepharose beads (GE Healthcare) for 2 hours. The beads were then washed five times with a lysis buffer to remove the immunocaptured proteins, boiled, and then subjected to western blot analysis using anti-Prdx5 (1:500, Ab Frontier) and anti-hnRNPK (1:500, Cell Signaling Technology) antibodies.

### LC-MS/MS analysis

The tryptic peptides were analyzed according to a modified method previously used for LC-MS/MS analysis (Lee et al., 2016). Briefly, the tryptic peptides were loaded onto an MGU-30 C18 trapping column (LC Packings, Amsterdam, The Netherlands). Concentrated tryptic peptides were eluted from the column and directed into a 10 cm × 75 μm I.D. C18 reverse phase column at a flow rate of 300 nL/min. The peptides underwent gradient elution in 0–55% acetonitrile over 100 min. MS and MS/MS spectra were acquired in the data-dependent mode using the LTQ-Velos ESI ion trap mass spectrometer (Thermo Fisher Scientific). For protein identification, MS/MS spectra were analyzed with MASCOT version 2.4 (Matrix Science, UK) using the mouse protein database downloaded from Uniprot. The mass tolerance for the parent or fragmentation was 0.8 Da. Carbamidomethylation of cysteine and oxidation of methionine were considered in MS/MS analysis as variable modifications of the tryptic peptides. The MS/MS data were filtered according to a false discovery rate criterion of 1%. Each sample was analyzed in triplicate. For protein quantification, we used the mol% value, which was calculated from the emPAI values in the MASCOT program (Lee et al., 2016; Yun et al., 2018). The canonical pathway of Prdx5-interacting proteins was screened using Ingenuity Pathway Analysis (IPA, Ingenuity Systems, Redwood City, CA, www.ingenuity.com), which leverages the Ingenuity Knowledge Base. Protein-protein interactions were constructed using STRING v11 (Szklarczyk et al., 2019).

### Luciferase reporter assays

MC3T3-E1 cells were cultured in α-MEM containing 10% FBS and 1% penicillin-streptomycin and transiently transfected with pGL3-OG2-Luc reporters using Lipofectamine 3000 (Invitrogen). The transfection efficiency was determined by co-transfecting the cells with a beta-galactosidase reporter (SV-β-gal). The reporter vectors were obtained from professor Won-Gu Jang, Daegu University, South Korea. The cells were transfected again with scrambled siRNA, Prdx5 siRNA, or pCMV-HA-Prdx5 plasmids. After the cells were recovered, osteoblast differentiation was induced by incubating them with 200 ng/mL BMP2 for 72 hours. Luciferase activity was measured using a luciferase reporter assay system (Promega) and a luminometer (SpectraMax i3x, Molecular Devices) according to the manufacturer’s instructions. The experiments were performed in triplicate and repeated thrice.

### RNA-seq analysis

BMMs were cultured for 4 days in an M-CSF and RANKL-containing medium for differentiating them into osteoclasts, and then lysed for RNA extraction. RNA was isolated using the RNeasy Mini Kit (Qiagen, Hilden, Germany), and quality control and sequencing were performed by Macrogen Inc. (Seoul, Republic of Korea). Briefly, a cDNA library was prepared using the TruSeq Stranded mRNA LT Sample prep kit (Illumina Inc.), and cDNA was synthesized using SuperScript II reverse transcriptase (Thermo Fisher Scientific).

All raw sequence reads were preprocessed using Trimmomatic (version 0.39) (Bolger et al., 2014) to remove adapter sequences and bases with low sequencing quality. The remaining clean reads were mapped based on the mouse reference genome (mm10) using Hisat2 (v2.1.0) (Kim et al., 2015) with the default parameters. BAM files generated by HiSat2 were further processed with Cufflinks (v2.2.1) (Trapnell et al., 2012) to quantify transcript abundance using the fragment per kilobase of exon per million fragments mapped (FPKM) normalization. Differential expression was analyzed using Cuffdiff (v2.2.1) to identify DEGs with FPKM > 1 in at least one sample and q-value < 0.05. We performed enrichment analysis of GO categories using the DAVID functional annotation tool (https://www.david.ncifcrf.gov). The mouse reference genome sequence and annotation data were downloaded from the UCSC genome browser (https://www.genome.ucsc.edu), and the R software was used to visualize the results.

### Statistics

Each experiment with cells was repeated at least thrice. Data are presented as mean ± standard error of the mean (SEM), unless indicated with the standard deviation (SD). The statistical analysis tests performed were a two-tailed Student’s *t*-test or one-way analysis of variance (ANOVA), followed by the least significant difference test for data with a normal distribution or the Kruskal–Wallis test for data not normally distributed. Image-based data were analyzed using the GraphPad Prism statistical software. Differences were considered statistically significant at \**p* < 0.05 and \*\**p* < 0.01.

### Data Availability

Proteomics data that support the findings of the current study have been deposited to the ProteomeXchange Consortium via the PRIDE (Perez-Riverol et al., 2019) partner repository with the dataset identifiers PXD020082 and 10.6019/PXD020082.

## Acknowledgements

This work was supported by the Korea Mouse Phenotyping Project (2014M3A9D5A0107365) of the Ministry of Science and ICT through the National Research Foundation.

---

**Figure 2–figure supplement 1.**
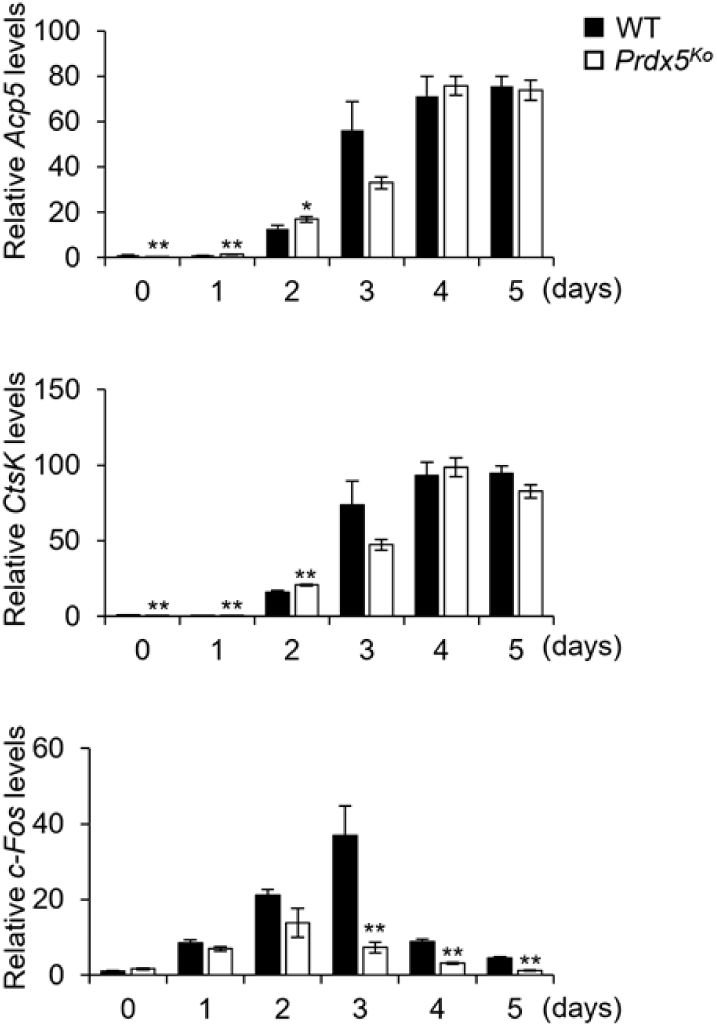
qRT-PCR was performed to determine the expression of osteoclast-related genes during osteoclastogenesis.

**Figure 2–figure supplement 2.**
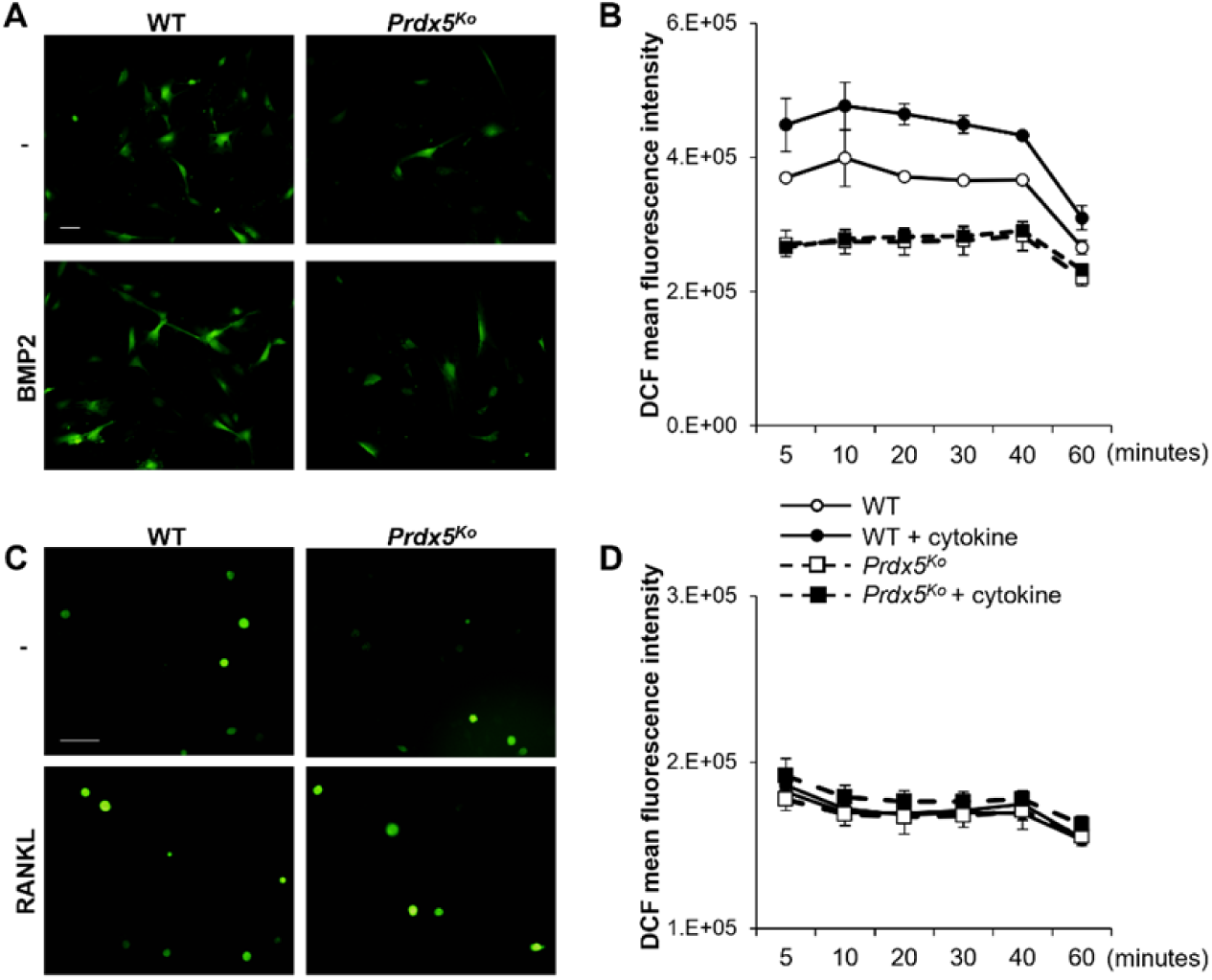
ROS levels are not altered in *Prdx5*-deficient osteoblasts. (A) Cellular ROS levels were measured via DCF fluorescence. The images were captured after 20 min of BMP2 stimulation of osteoblasts. Scale bar, 100 µm. (B) ROS levels were measured at indicated time periods after BMP2 stimulation. (C) Cellular ROS levels were measured after RANKL stimulation in osteoclasts. Scale bar, 100 µm. (D) ROS levels were measured at indicated times after RANKL stimulation.

**Figure 3–figure supplement 1.**
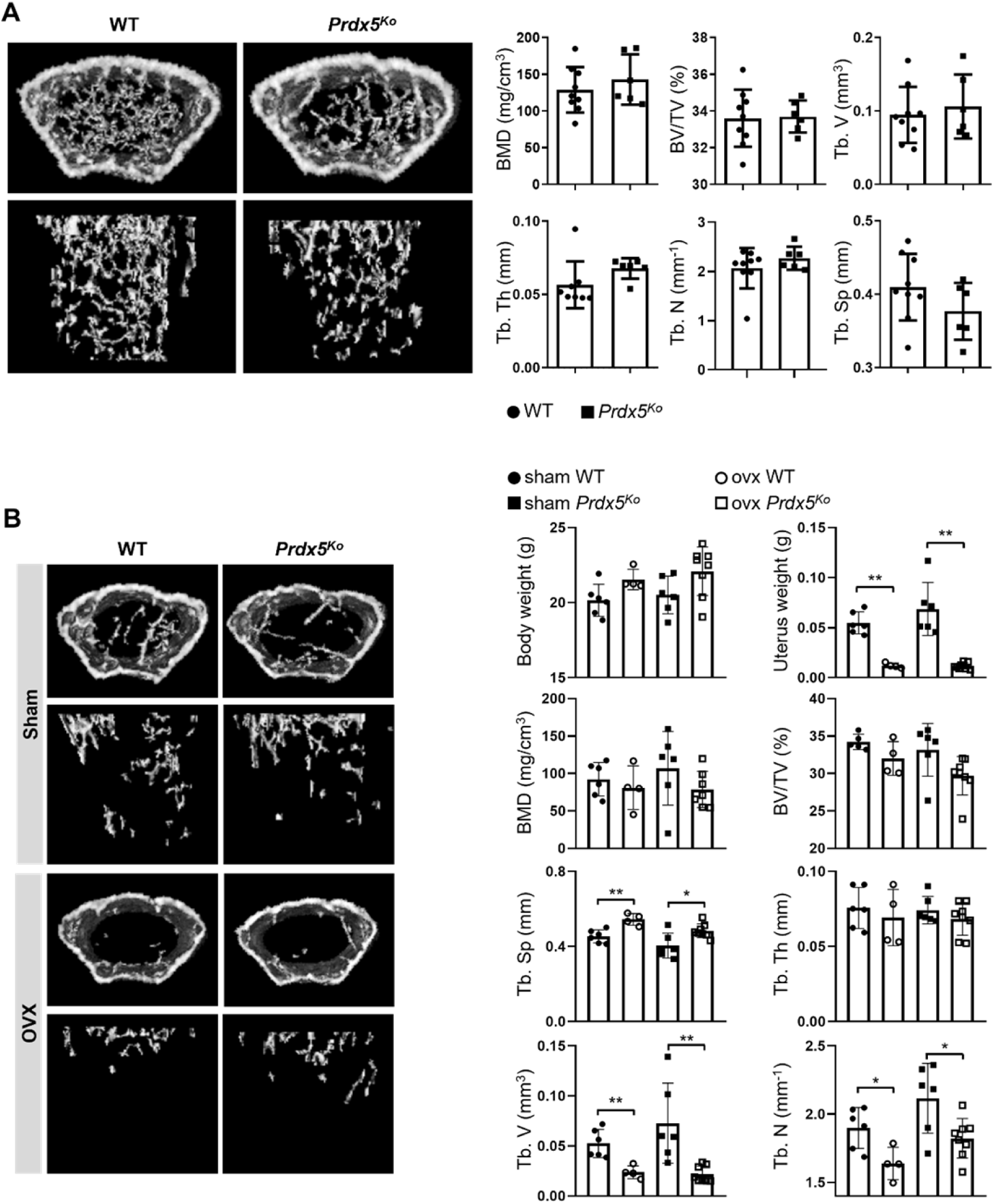
Female mice show normal phenotypes. (A) Micro-CT images of femurs from WT and *Prdx5*^Ko^ mice at 12 weeks. Micro-CT data were quantified (n = 6**–**9). BMD, bone mineral density; BV/TV, bone volume relative to total tissue volume; Tb. V, trabecular volume; Tb. Th, trabecular bone thickness; Tb. N, trabecular bone number; Tb. Sp, trabecular bone space. (B) OVX or sham surgery was performed on 10-week-old females that were sacrificed after 4 weeks for micro-CT analysis (n = 4**–**8)

**Figure 4–figure supplement 1.**
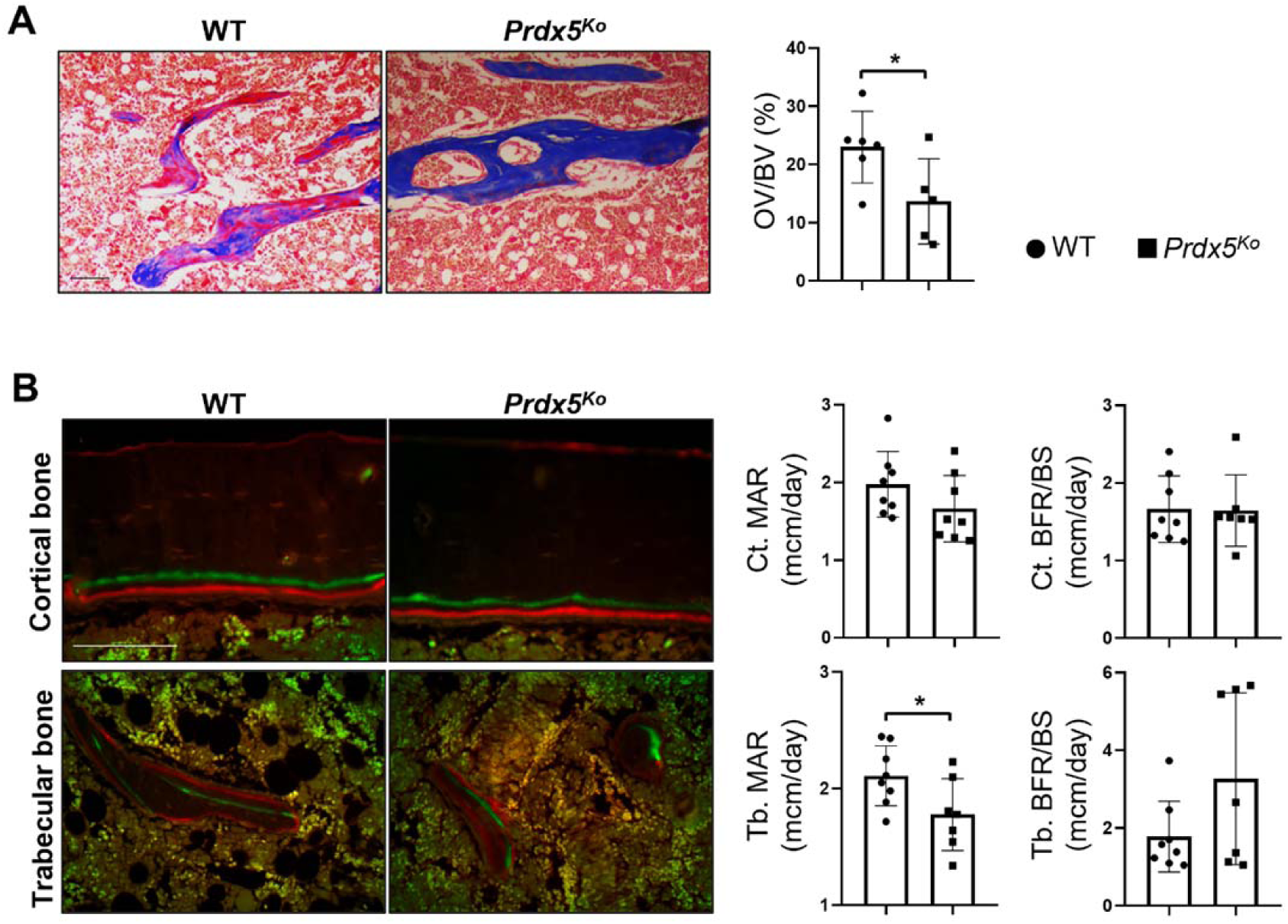
*Prdx5*^Ko^ mice show reduced bone turnover. (A) Differential staining of the bone (blue) and osteoid (red in bone) was performed using Goldner’s trichrome method. The ratio of osteoid volume/bone volume (OV/BV) was measured using the Bioquant Osteo program. (B) Bone turnover parameters of the femurs from 8-week-old mice were measured via dynamic bone histomorphometry after serial injections of calcein and alizarin red S. Two-color labeled mineralized fronts, visualized via fluorescence micrography, indicated a low bone turnover with reduced MAR in the trabecular bone, but not in the cortical areas of *Prdx5^ko^* compared to that in WT mice. MAR: mineral apposition rate. BFR/BS: bone formation rate per bone surface. (n = 5**–**7). Data are presented as mean ± SEM.

**Figure 5–figure supplement 1.**
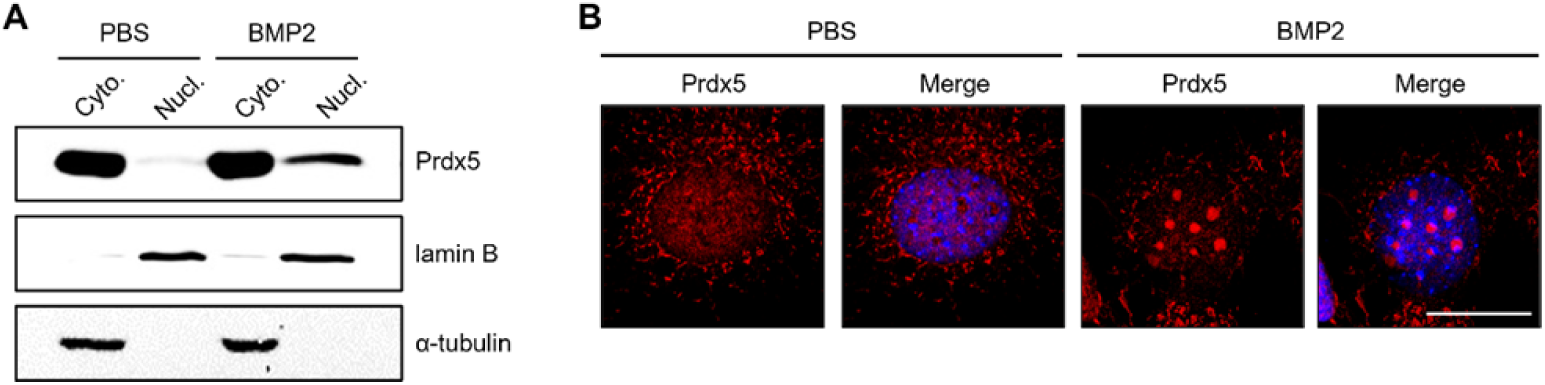
BMP2 induces nuclear translocation of Prdx5. (A) Western blot analysis of Prdx5 in the cytoplasmic and nuclear fractions of osteoblasts treated with BMP2 for 4 days. (B) Fixed osteoblasts after 4 days of BMP2 treatment were stained with an anti-Prdx5 antibody (red) and imaged using confocal microscopy. The nucleus was counterstained in blue. Scale bar, 20 μM

**Figure 7–table supplement 1.**
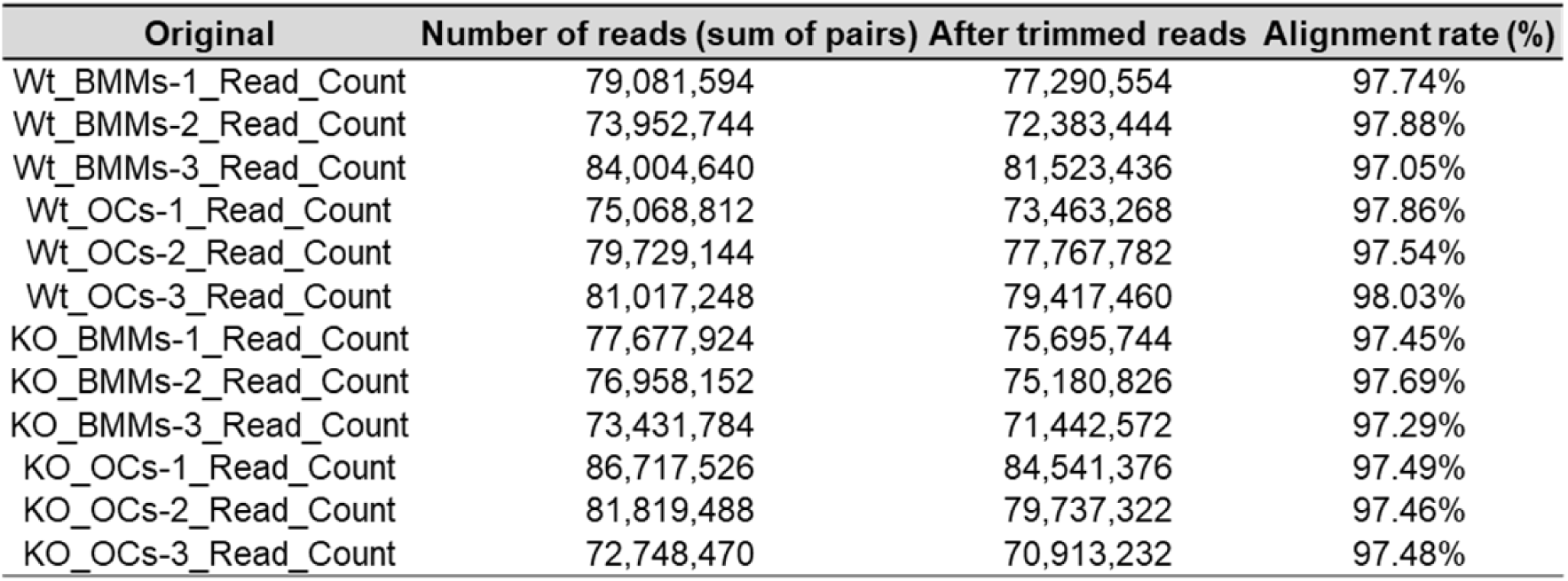
Statistics of RAN-seq analysis

**Figure 7–figure supplement 1.**
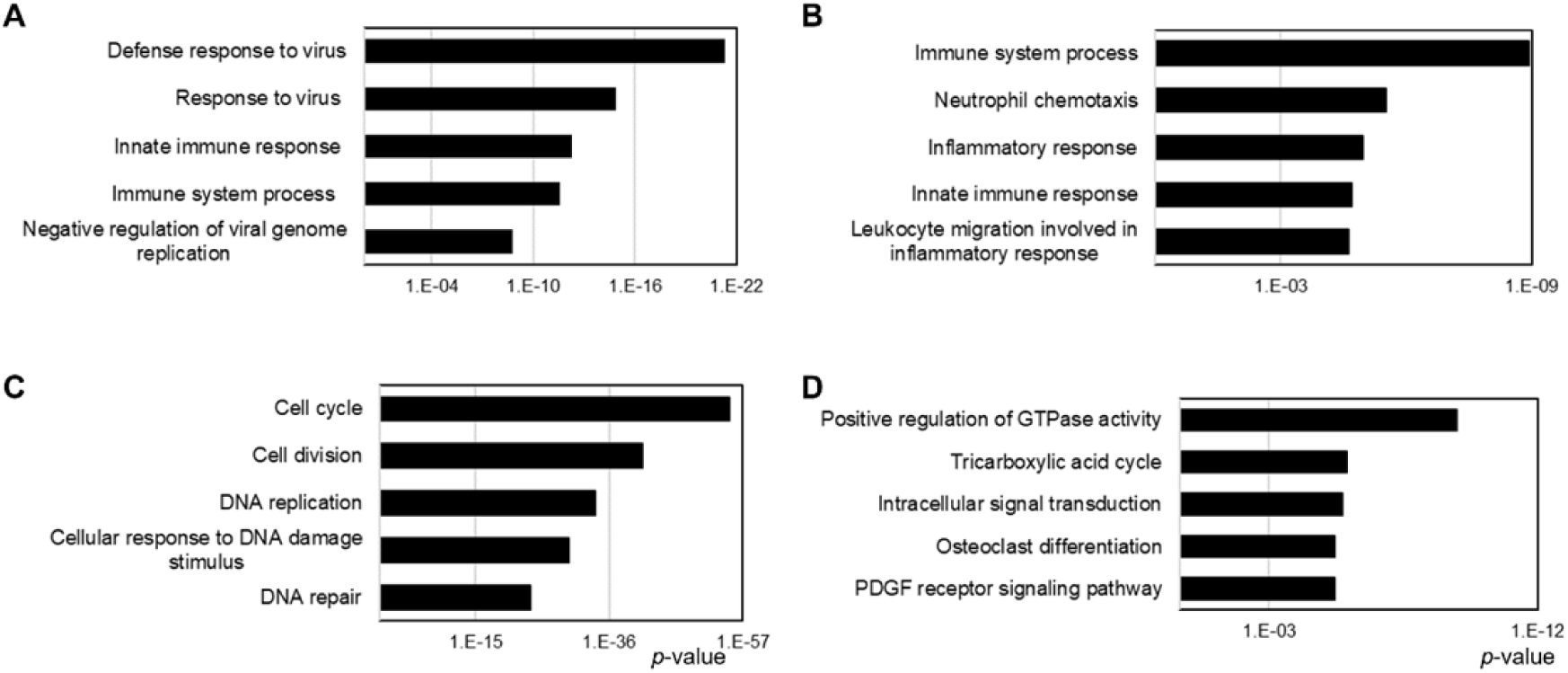
GO analysis of DEGs via RNA-seq analysis. (A) Total 153 downregulated and (B) 61 upregulated DEGs in BMMs. (C) Total 616 downregulated and (D) 641 upregulated DEGs in OCs. Top five biological pathways are represented by the *p*-value (X-axis).

